# 5’tRNA-derived fragments modulate β-cell homeostasis and islet macrophage activation in type 2 diabetes

**DOI:** 10.1101/2025.07.22.665911

**Authors:** Cristina Cosentino, Rémy Klein, Véronique Menoud, Claudiane Guay, Elena Aiello, Stefano Auddino, Gianfranco Di Giuseppe, Gea Ciccarelli, Alessandra Galli, Francesco Alabiso, Eleonora Mangano, Flora Brozzi, Karim Bouzakri, Stefania D’Adamo, Silvia Cetrullo, Giuseppe Quero, Andrea Mari, Sergio Alfieri, Andrea Giaccari, Teresa Mezza, Francesco Dotta, Guido Sebastiani, Romano Regazzi

**Affiliations:** Department of Fundamental Neuroscience, University of Lausanne, Lausanne, Switzerland; Dipartimento di Scienze Mediche, Chirurgiche e Neuroscienze, University of Siena, Italy; Endocrinology and Diabetology Unit, University Hospital Agostino Gemelli, Rome, Italy; Department of Translational Medicine and Surgery, Catholic University of Sacred Heart, Rome, Italy; Department of Biomedical and Neuromotor Sciences, University of Bologna, Bologna, Italy; C.N.R. Institute for Biomedical Technologies, Milan, Italy; UMR DIATHEC, EA 7294, Centre Européen d’Etude du Diabète, Université de Strasbourg, Fédération de Médecine Translationnelle de Strasbourg, Strasbourg, France; Digestive Surgery Unit, University Hospital Agostino Gemelli, Rome, Italy; Institute of Neuroscience, National Research Council, Padua, Italy; Department of Biomedical Sciences, University of Lausanne, Lausanne, Switzerland

## Abstract

During obesity and type 2 diabetes, pancreatic β-cells face chronic environmental stress, while islet-resident macrophages (iMACs) undergo metabolic reprogramming that exacerbates β-cell dysfunction. Stress-induced cleavage of transfer RNAs (tRNAs) generates tRNA-derived fragments (tRFs), whose role in this context is not fully understood. We identify elevated levels of 5’tRF^Glu(CTC)^ and 5’tRF^Gly(GCC)^ in β-cells and iMACs from db/db mice and in islets from type 2 diabetic patients. Notably, 5’tRF^Glu(CTC)^ is also induced under prediabetic conditions and inversely correlates with insulin secretion. Lipotoxic stress triggers their production via Angiogenin-mediated cleavage. Blocking 5’tRF^Glu(CTC)^ in islets protects against β-cell apoptosis and restores insulin secretion under palmitate stress. Using a β-cell/macrophage co-culture system, we show that β-cell contact shapes a unique macrophage phenotype (iMAC-like) that shifts upon palmitate exposure—recapitulating in vivo observations. Inhibiting 5’tRF^Glu(CTC)^ in iMAC-like cells prevents this activation switch, reduces β-cell stress, and improves insulin secretion. Mechanistically, 5’tRF^Glu(CTC)^ interacts with RNA-binding proteins to regulate transcriptional and post-transcriptional pathways linked to immune activation, extracellular matrex remodeling, neurogenesis, and oxidative stress. Our study identifies 5’tRFs as key mediators of islet microenvironment remodeling in diabetes, offering new insights into intercellular stress signaling in metabolic disease.

## INTRODUCTION

Obesity, characterized by excessive caloric intake and increased adiposity, represents a significant risk factor for type 2 diabetes (T2D) development. Pancreatic β-cells play a pivotal role in maintaining glucose homeostasis by secreting insulin in response to rising blood glucose levels. However, inappropriate nutritional conditions trigger endoplasmic reticulum (ER) stress and oxidative stress, which collectively drive β-cell dysfunction and, eventually, apoptosis. Simultaneously, the energy imbalance established during obesity activates innate immune responses as an adaptive mechanism to cope with the increased nutrient load. Indeed, expansion of islet macrophages (Mφs) was observed in T2D patients and in rodent models of obesity and T2D^1–4^. Under physiological conditions, islet resident Mφs (iMACs) exhibit a unique transcriptional profile characterized by the expression of pro-inflammatory markers, such as IL-1β, that sustain proper β-cell function^5^. However, high fat diet (HFD) induces iMAC expansion accompanied by profound transcriptional remodeling: while certain genes associated with the pro-inflammatory profile are upregulated, others are notably downregulated^3^. In addition to these changes, β-cell death was recently reported to activate a reparative and anti-inflammatory state in Mφs, marked by the secretion of Insulin-like Growth Factor 1 (IGF1), which enhances β-cell function^6^. These findings underscore the dynamic and multifaceted role of Mφs in islet health and T2D pathogenesis.

Despite significant progress in understanding islet physiopathology, the molecular mechanisms that regulate β-cell stress responses and the metabolic reprogramming of islet-resident macrophages (iMACs) during obesity remain poorly defined. In particular, the signals mediating the crosstalk between β-cells and iMACs are still largely unknown.

Recent genome-wide association studies and high-throughput sequencing approaches have pointed to a potentially crucial role of transfer RNA (tRNA) dynamics in maintaining pancreatic β-cell homeostasis. Traditionally known for their role in protein synthesis, tRNAs are now recognized as being tightly regulated by both nutritional and environmental cues. Post-transcriptional modifications and enzymatic cleavage of tRNAs have been linked to cellular stress responses and the development of various diseases^7, 8^.

tRNA cleavage gives rise to tRNA-derived fragments (tRFs), a novel class of small non-coding RNAs that can influence numerous cellular functions, including gene expression, mRNA translation, cell differentiation, and apoptosis^9^.

Notably, loss-of-function mutations in the tRNA-modifying enzyme TRMT10A have been associated with a syndrome marked by microcephaly and early-onset diabetes. Impaired TRMT10A function leads to altered tRF biogenesis, contributing to β-cell oxidative stress and apoptosis^10, 11^. Moreover, we have previously shown that tRFs are dynamically regulated during β-cell maturation^12^ and in the pancreatic islets of rodent models predisposed to diabetes^13^.

Despite these insights, the functional role and mechanisms of action of tRFs in β-cells remain unclear. In addition, their regulation and function in iMACs and in the intercellular communication between iMACs and β-cells during obesity has not been investigated so far.

This study provides evidence that obesity and diabetes induce a specific modulation of tRFs in both β-cells and iMACs. Indeed, 5’tRFs derived from tRNA^Glu(CTC)^ and tRNA^Gly(GCC)^ were found to be induced in both cell types, an effect recapitulated *in vitro* by exposure to saturated fatty acids. Blockade of 5’tRF^Glu(CTC)^ in islet cells or specifically in macrophages had protective effects against lipotoxicity. The analysis of the 5’tRF^Glu(CTC)^ interactome revealed an association with RNA-binding proteins involved in mRNA processing, splicing and translation. In agreement with this finding, blockade of this tRNA fragment induced transcriptional and translational changes.

## RESULTS

### Obesity and type 2 diabetes induce changes in 5’tRFs in β-cells and iMACs

To investigate whether tRNA cleavage and consequently the pool of tRFs is affected by obesity and diabetes, we isolated the pancreatic islets of 8-week-old overweight and hyperglycemic db/db mice and of age-matched lean and normoglycemic wild type controls (**Figure S1.a-b**). Db/db mice display hyperphagia due to a mutation in the leptin receptor gene, and develop obesity and diabetes. It was previously shown that at 8 weeks of age, in the initial phases of the disease, islet macrophages (iMAC) of db/db mice expand and undergo transcriptional remodeling^6^. Since both iMACs and β-cells are highly sensitive to nutritional stress and to the diabetogenic microenvironment, we purified the two cell populations by FACS (**Figure 1a; Figure S1.c**). Although the effect was not statistically significant, the number of isolated iMACs tended to increase in db/db mouse islets compared to those of wild type mice (**Figure S1.d**). Characterization of the tRF profile of FACS sorted cells led to the identification of 152 tRFs displaying significant changes in β-cells and 21 in iMACs of db/db mice compared to wild type controls (FC>2, FDR p<0.1) (**Figure 1b and d**). After removal of redundant sequences with differences of 1-2 nucleotides at the cleavage site, we identified a group of tRFs derived from 17 cytosolic tRNA isoacceptors in β-cells and 11 in iMACs being modulated in db/db mice (**Figure 1.c and e)**. Three tRFs derived from the 5’end of tRNA^Glu(CTC)^, tRNA^Gly(GCC)^ and tRNA^Asp(CTC)^ were induced in both cell types. The upregulation of 5’tRF^Glu(CTC)^ and 5’tRF^Gly(GCC)^ was validated by qPCR (**Figure 1.f1-4**), while qPCR for 5’tRF^Asp(CTC)^ was not conclusive due to the presence of multiple melting curves. Interestingly, we previously demonstrated that 5’tRF^Glu(CTC)^ is repressed during islet cell maturation and may regulate newborn β-cell function^12^.

**Figure 1.**
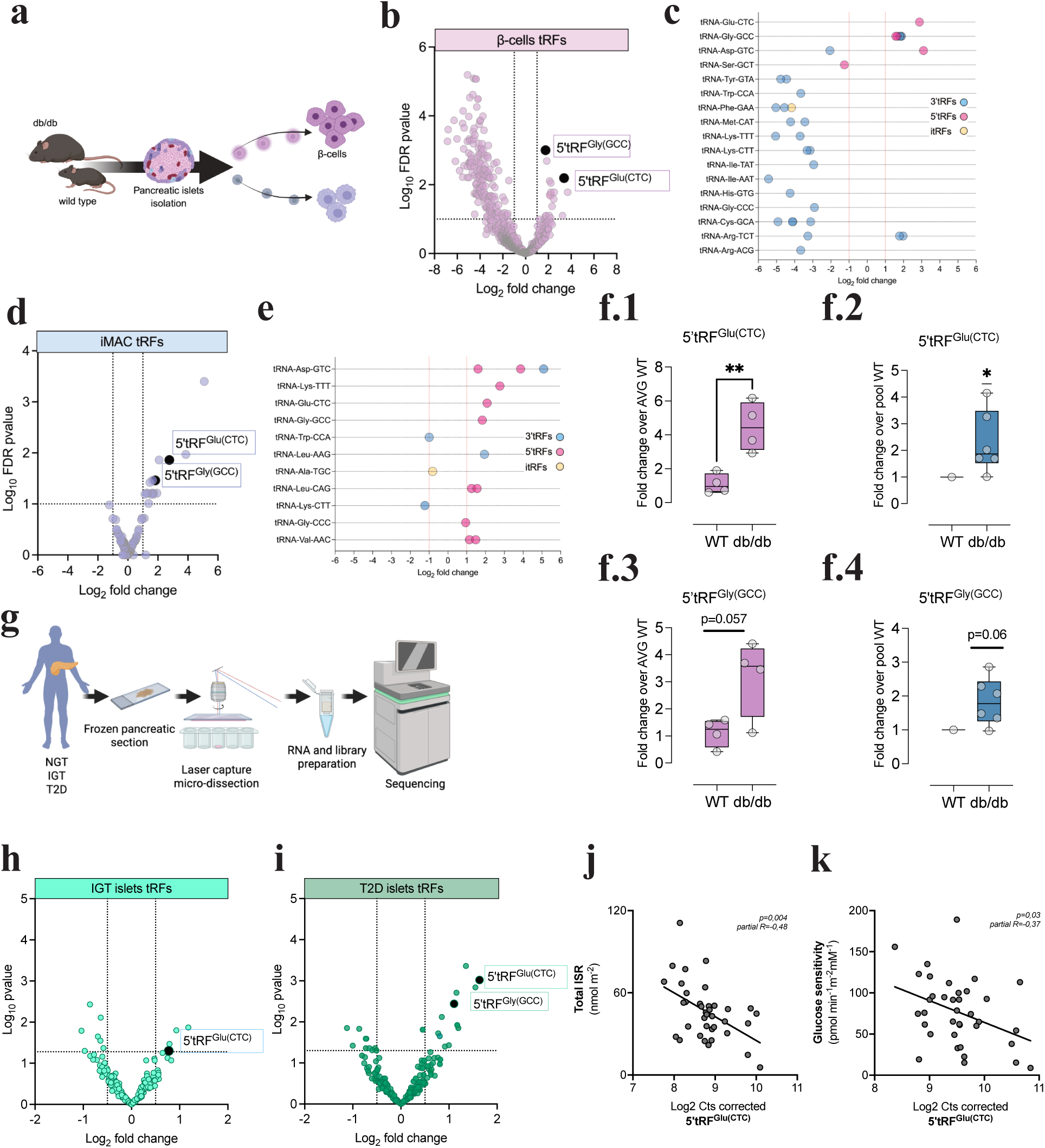
tRF profile of pancreatic islet cells in obesity and T2D: Pancreatic islets were obtained from db/db mice and wild type controls; *β*-cells and iMACs were isolated by FACS (**a**) and small RNA sequencing was performed (**b-e**). Volcano plots show differentially expressed tRFs in db/db *β*-cells (**b**) and iMAC (**d**) compared to controls. tRNA isodecoder of origin and the different types of tRFs are described in (**c)** and (**e**). 5’tRF^Glu(CTC)^ and 5’tRF^Gly(GCC)^ levels were detected by qPCR and normalized with the expression of Let-7e in *β*-cells (**f1-f3**) or miR-16 in iMAC (**f2-f4**) (*p<0.05, **p<0.01 by Student t-test or Wilcoxon signed rank test). Human islets were collected by laser capture microdissection from pancreatic sections (**g**). Patients were categorized based on oral glucose tolerance: normal glucose tolerance (NGT, n=12 <140 mg/dL 2 hours plasma glucose); impaired glucose tolerance (IGT n=13 140-199 mg/dL 2 hours plasma glucose) and type 2 diabetes (T2D n=11 ≥200 mg/dL 2 hours plasma glucose). Volcano plots recapitulate the changes in tRF levels in IGT versus NGT islets (**h**) and T2D versus NGT islets (**i**). The association of 5’tRF^Glu(CTC)^ levels with the clinical parameters total insulin secretion rate (ISR) (**j**) and glucose sensitivity (**k**) was performed using linear models. The association of Log2 scaled tRF expression values with clinical/metabolic parameters was corrected for the covariates (Age, Gender, BMI).

To analyze the modulation of tRFs in the islets of human living donors, we collected RNA from islets isolated by laser capture microdissection (LCM) from n=12 normal glucose tolerant (NGT), n=13 impaired glucose tolerant (IGT) and n=11 T2D diagnosed patients (**Figure 1.g**; **Table S1**). Small RNA sequencing and tRF profiling identified 8 tRFs significantly changed in the IGT group (**Figure 1.h**) and 13 tRFs modulated in the T2D (**Figure 1.i**) compared to the NGT controls (FC>1.5, p<0.05). We found that the 5’tRF^Glu(CTC)^ was significantly upregulated in both IGT and T2D groups (**Figure 1.h-i**), while 5tRF^Gly(GCC)^ was increased only in T2D islets (**Figure 1.i**). Notably, these tRFs present 100% of sequence homology between mouse and human. Our findings suggest that tRF modulation in islet cells occurs during T2D development and is conserved between species.

Association studies revealed that islet 5’tRF^Glu(CTC)^ levels were negatively associated to basal and total insulin secretion rates and glucose sensitivity (**Figure 1.j-k, Table S2**). Notably, this association was mainly driven by the IGT group (**Figure S1.e**). In contrast, no correlations were found with basal or mean glucose levels during the OGTT (**Table S2**). These findings suggest that islet 5’tRF^Glu(CTC)^ levels may directly impair in vivo insulin secretory capacity and reflect early decline of β-cell function, independently of both glucose control and glucose tolerance status.

### 5’tRF^Glu(CTC)^ and tRF^Gly(GCC)^ are induced by exposure to saturated free fatty acids

We next investigated whether *in vitro* exposure to fatty acids—mimicking the lipotoxic stress associated with obesity-linked diabetes— recapitulates the induction of the tRFs observed *in vivo*. The unsaturated fatty acid oleate did not modulate the tRF levels in MIN6 β-cells (**Figure 2.a**). We observed a significant increase of 5’tRF^Glu(CTC)^ induced by the saturated free fatty acids stearate and palmitate (**Figure 2.a**). Interestingly, palmitate elicited a greater rise of 5’tRF^Glu(CTC)^ and induced also the 5’tRF^Gly(GCC)^ levels compared to the less lipotoxic stearate (**Figure 2.a**). This effect appeared to be time dependent (**Figure S2.a**). Palmitate-dependent tRF induction was confirmed in primary islets derived from wild type mice and from human cadaveric donors (**Figure 2.b-c**). qPCR analysis of FACS-sorted β-cells and iMACs confirmed the induction of the tRFs in both cell types upon 48h treatment with palmitate (**Figure 2.d-e**).

**Figure 2.**
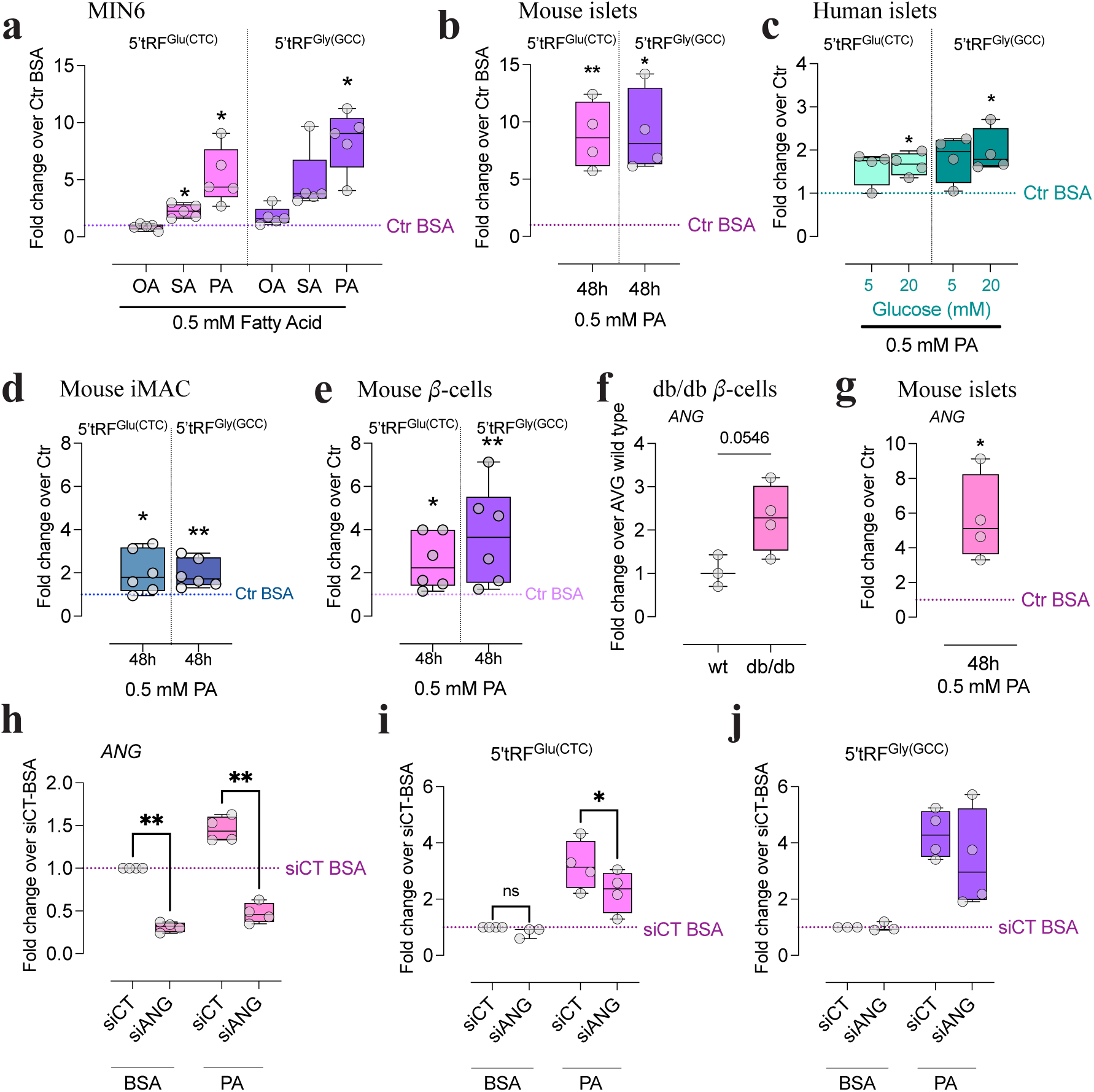
5’tRFs from tRNA^Glu(CTC)^ and tRNA^Gly(GCC)^ are induced by fatty acids in *β*-cells and iMACs: MIN6 cells were treated with 0.5 mM of the unsaturated fatty acid oleate (OA) or the saturated fatty acids stearate (SA) and palmitate (PA) for 48 hours and the levels of 5’tRFs were measured by qPCR; 5’tRFs levels were normalized against basal 0.75% BSA treatment condition (**a**). The induction of 5’tRF^Glu(CTC)^ and 5’tRF^Gly(GCC)^ was assessed in dispersed mouse islets treated with PA (**b**) and human islets treated with PA alone or in combination with 20mM Glucose for 48h (**c**). Upon exposure to palmitate, mouse islets were dispersed, iMACs and *β*-cells were isolated by FACS and tRFs were measured in the two cell populations (**d-e**). Angiogenin (ANG) expression was assessed by qPCR in FACS sorted *β*-cells from db/db and wild type (wt) mice (**f**) and dispersed mouse islets treated with PA (**g**). Knock down of Angiogenin was achieved by siRNA transfection (siANG) in MIN6 cells (**h**). The levels of 5’tRF^Glu(CTC)^ and 5’tRF^Gly(GCC)^ upon Angiogenin knock down was assessed by qPCR (**i-j**). *p<0.05, **p<0.01 by one-way ANOVA with Sidak correction.

In mouse and human islet cells, 5’tRF^Glu(CTC)^ fragments are 29-30nt long (**Table S3**) and are generated by a cleavage adjacent or in the anticodon loop, consistent with a classification as tRNA halves. Angiogenin was previously identified as the major enzyme responsible of tRNA cleavage at the level of the anticodon loop and of the biogenesis of tRF-halves^14, 15^. 5’tRF^Gly(GCC)^ fragments identified in our datasets are 26-30nt long (**Table S3**). The biogenesis of these 5’tRFs have been also associated with Angiogenin expression, although other RNAses may participate to the trimming of the cleavage site^14^. We have previously showed that silencing of Angiogenin gene (*ANG*) in islets from newborn rats decreases the levels of 5’tRF^Glu(CTC)12^. To assess whether this enzyme is responsible for the generation of 5’tRF^Glu(CTC)^ and 5’tRF^Gly(GCC)^ under obesity and diabetes conditions, we first measured the expression of Angiogenin in our experimental models. We observed that *ANG* mRNA is increased in β-cells of db/db mice compared to wild type controls (**Figure 2.f**) and is induced in MIN6 and dispersed mouse islet cells upon palmitate exposure (**Figure S2.b and 2.g**). To further confirm the role of Angiogenin in 5’tRF^Glu(CTC)^ and 5’tRF^Gly(GCC)^ biogenesis, we decreased the level of the enzyme using a siRNA (**Figure 2.h**). We found that transient knockdown of Angiogenin attenuates the induction of 5’tRF^Glu(CTC)^ but not of 5’tRF^Gly(GCC)^ (**Figure 2.i-j**). These findings validate our previous observations in newborn β-cells and suggest that additional enzymes may be involved in palmitate-dependent tRNA cleavage.

### Blockade of 5’tRF^Glu(CTC)^ attenuates lipotoxic damage in β-cells

Next, we investigated the impact of a global reduction of 5’tRF^Glu(CTC)^ in islet cells on β-cell homeostasis. For this purpose, we used an antisense oligonucleotide (ASO)-based approach to target and inhibit 5’tRF^Glu(CTC)^ (α*-*Glu) under basal and palmitate conditions (**Figure 3.a**). Transfection of α*-*Glu ASO resulted in >90% reduction of 5’tRF^Glu(CTC)^ in mouse islet cells (**Figure 3.b**) and 70-90% reduction in MIN6-β-cells (**Figure S.2c**). Notably, α*-*Glu ASO did not affect full length tRNA^Glu(CTC)^ levels or its aminoacylation (**Figure S2.d-e**). We observed that the inhibition of 5’tRF^Glu(CTC)^ in mouse islet cells prevents palmitate-induced β-cell apoptosis measured by Caspase 3 cleavage (**Figure 3.c-d**). Next, we assessed the consequence of 5’tRF^Glu(CTC)^ blockade on insulin secretion in MIN6-β-cells. Upon palmitate treatment, insulin release in response to glucose, expressedas the percentage of the insulin content, was reduced (**Figure 3.e**). This effect was ameliorated by the inhibition of 5’tRF^Glu(CTC)^. As expected, prolonged palmitate treatment led also to an important decrease in insulin content (**Figure 3.f**). Consequently the amount of insulin released was strongly diminished when normalized for total protein content (**Figure 3.g**). This effect was not prevented by α*-*Glu transfection.

**Figure 3.**
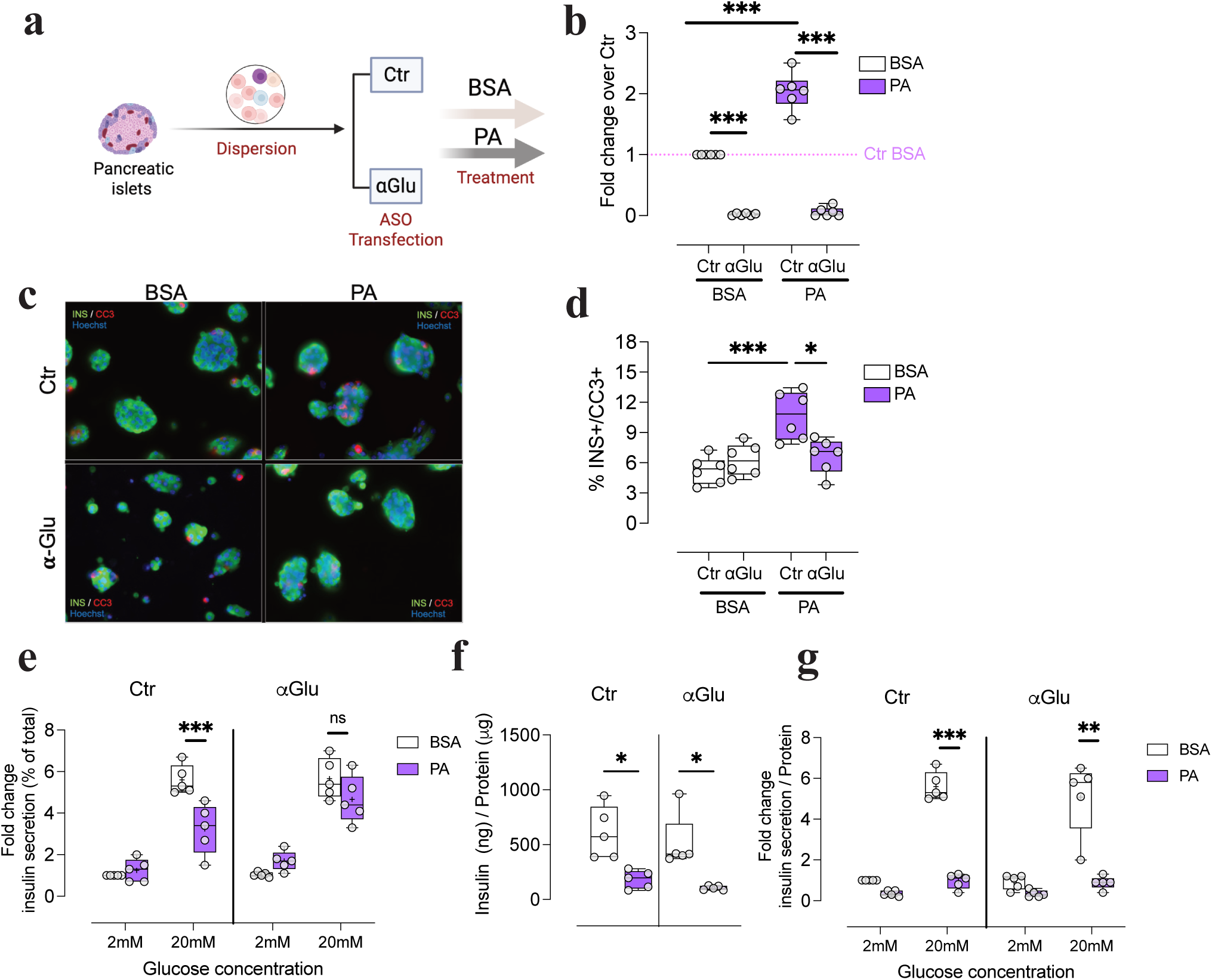
5’tRF^Glu(CTC)^ inhibition in islet cells protects against lipotoxicity: dispersed mouse islet cells were transfected with an antisense oligonucleotide (ASO) targeting 5’tRF^Glu(CTC)^ (**α**GLU) or an antisense of control (Ctr) and were exposed to either vehicle (BSA) or palmitate (PA) for 48 hours (**a**). The levels of 5’tRF^Glu(CTC)^ were assessed by qPCR and normalised by the geometric mean of Let7a and miR-7 expression (**b**). Insulin (green) and cleaved caspase (red) positive cells were detected by immunofluorescence (**c**) and the percentage of double positive over total insulin positive cells was calculated (**d**). Cellular insulin content and released insulin was measured by ELISA (**e-g**) in BSA or PA treatment. Glucose-induced insulin secretion was estimated in basal (2mM glucose) and stimulating (20mM glucose) conditions and expressed as percentage of total insulin content (**e**) or normalized by total protein content (**g)** or . *p<0.05, **p<0.01, ***p<0.001, ****p<0.0001 by One-way ANOVA with Sidak multiple comparison correction (a,c and d) and by Three-way ANOVA with Tukey correction (e and f).

### Blockade of 5’tRF^Glu(CTC)^ in macrophages ameliorates β-cell homeostasis upon palmitate exposure

In addition to endogenous processes disrupting β-cell homeostasis, the proliferation and activation of iMACs is expected to play a critical role in the development of diabetes under obesity conditions. We asked whether the modulation of 5’tRF^Glu(CTC)^ specifically in macrophages contributes to these pathogenic mechanisms. To this end, we aimed to target and inhibit 5’tRF^Glu(CTC)^ in iMACs and assess the resulting effects on both Mφ phenotype and β-cell homeostasis. However, manipulating primary iMACs poses significant challenges, as these cells, once isolated by FACS, rapidly lose their phenotype and are not viable when cultured alone. To overcome these limitations and since the activation state of iMACs is mainly determined by the crosstalk with surrounding endocrine cells, we established an *in vitro* model to explore these interactions under controlled conditions. To this end, bone marrow-derived cells from wild type mice were differentiated *in vitro* into naïve (M0) Mφs (**Figure S3.a**). M0 Mφs were further differentiated by culturing them in direct contact with MIN6 β-cells to induce an iMAC-like phenotype (**Figure 4a**). After 3 days of co-culture, the cells were separated by FACS (**Figure S3.b**). The iMAC-like phenotype was evaluated by the expression of gene markers for Mφ polarization. Compared to M0 Mφs, iMAC-like cells showed an upregulation of the pro-inflammatory marker *IL-1*β and, at the same time, an induction of the anti-inflammatory markers *Mrc1* and *Ym1* (**Figure 4.b**). This mixed phenotype with high *IL-1*β expression recapitulates previous observations on the activation state of iMACs *in vivo*^16^. We next evaluated the impact of PA treatment and of the modulation of 5’tRF^Glu(CTC)^ on iMAC-like cells. To this purpose, M0 Mφs were transfected with either Ctr or αGlu ASO prior co-culture and then treated with BSA or PA (**Figure 4.c**). In FACS-sorted iMAC-like cells trasfected with Ctr oligonucleotide PA raised 5’tRF^Glu(CTC)^ levels, while αGlu transfection led to a decrease of the tRF in both BSA and PA conditions (**Figure 4.d**). We observed that PA decreased *IL-1*β expression in Ctr iMAC-like cells and upregulated the homeostatic genes *Mrc1* and *Ym1* (**Figure 4.e** – full bars). These results matched the transcriptomic modulation observed in iMACs from mice fed with high-fat diet^3^ and from db/db mice^6^. At the same time, we observed that 5’tRF^Glu(CTC)^ inhibition led to a increase of *IL-1*β expression in basal conditions and prevented the induction of *Mrc1* and *Ym1* elicited by PA exposure (**Figure 4.e** – pattern bars). These data suggest that the 5’tRF^Glu(CTC)^ may be a key modulator of iMAC activation switch in response to environmental stimuli.

**Figure 4.**
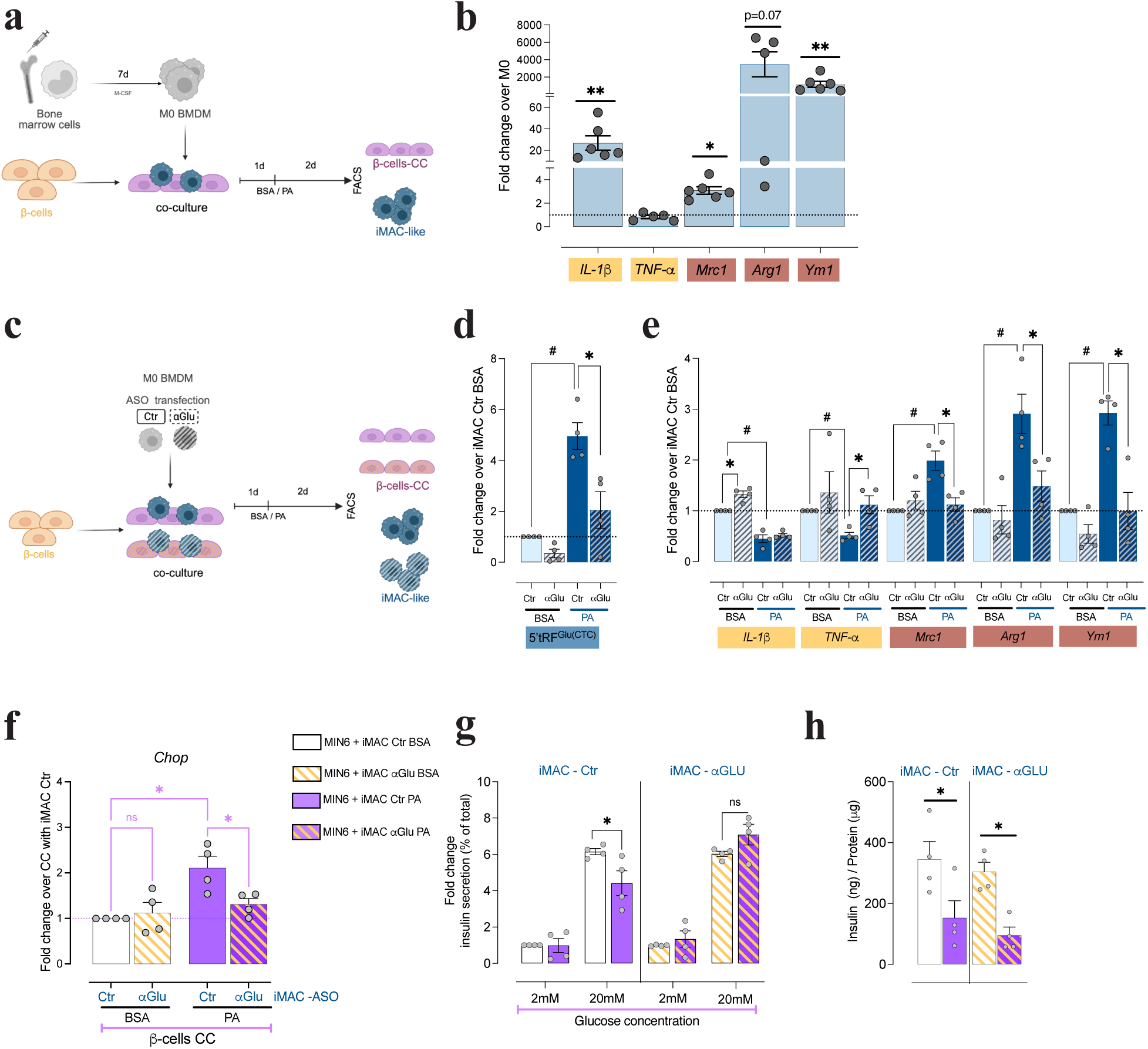
5’tRF^Glu(CTC)^ inhibition in iMAC-like cells attenuates the phenotypic switch induced by palmitate and protects co-cultured *β*-cells: schematic representation of co-culture experiment: bone marrow derived cells were differentiated for 7 days with M-CSF; naïve (M0) BMDMs were co-cultured with MIN6 *β*-cells for 3 days and the two cell populations were separated by FACS (**a**). The expression of polarization markers in iMAC-like cells was compared with M0 M*φ*s (**b**). Schematic representation of co-culture experiment in which M0 M*φ*s were previously transfected with ASO of Ctr or *⍺*GLU (**c**). 5’tRF^Glu(CTC)^ and polarization markers were measured in iMAC-like cells transfected with *⍺*GLU (pattern bars) or Ctr (full bars) in palmitate or BSA conditions (**d-e**). *Chop* expression (**f**), glucose-induced insulin secretion (**g**) and insulin content (**h**) were assessed in MIN6 *β*-cells cultured with iMAC-like cells of Ctr (full bars) or transfected with *⍺*GLU (pattern bars) in BSA or PA treatment conditions. *p<0.05, **p<0.01 ***p<0.001 *⍺*Glu versus Ctr; #p<0.05 ###p<0.001 PA versus BSA by One-way ANOVA with Sidak correction for multiple comparisons.

To assess whether the co-culture with Mφs modulates MIN6 β-cell responses to PA, we isolated the insulin-secreting cells by FACS and we measured the expression of *Chop* which is induced by ER-stress during lipotoxicity^17^. Compared to MIN6 cells cultured alone, we did not observe any significant modulation of this gene by the presence of non-transfected Mφs (**Figure S3.c**). However, co-culture with αGlu iMACs - lacking 5’tRF^Glu(CTC)^ – prevented *Chop* induction during PA treatment (**Figure 4.f**). While PA still decreased insulin content and secretion in MIN6 co-cultured with Ctr Mφs, the percentage of insulin content released in response to glucose was restored by the co-culture with iMACs depleted from 5’tRF^Glu(CTC)^ (**Figure 4.g-h**).

Taken together, our observations suggest that targeting 5’tRF^Glu(CTC)^ in Mφs may have indirect beneficial effects on β-cells, probably due to the attenuation of the phenotypic switch induced by saturated fatty acid exposure.

### 5’tRF^Glu(CTC)^ interacts with RNA-binding proteins and participates to gene expression regulation

We then explored the mechanism by which 5’tRF^Glu(CTC)^ exerts its regulatory effect in islet cells. tRFs can mediate a variety of cellular functions depending on the molecular interactions in which they are engaged. Therefore, we decided to identify the protein interactors of 5’tRF^Glu(CTC)^. A biotinylated tRF mimic (GLU) or a scrambled control (CTRL) was transfected in MIN6 cells (**Figure S4.a**); following UV-crosslink, cell lysates were pulled-down and analysed by mass spectrometry (**Figure 5.a**). This technique allowed the identification of 25 proteins likely interacting with 5’tRF^Glu(CTC)^ (**Figure 5.b**). The interactions with Msi2, the mostly enriched protein, and with hnRNP-A3, the most abundant protein in the 5’tRF^Glu(CTC)^ pull-down, were validated by western blot (**Figure 5.c**). Functional enrichment revealed an overrepresentation of GO terms “RNA splicing” and “RNA processing” in biological processes and “spliceosome complex”, “ribonucleoprotein granules” and “nuclear speckles” in cellular components (**Figure S4.c-d**). Indeed, 5’tRF^Glu(CTC)^ interactors consist in RNA-binding proteins (RBPs) involved in several steps of gene expression regulation, such as splicing, transport and translation (**Figure 5.d**). To further characterize the connectivity among these RBPs, we performed STRING-based protein-protein interaction analysis, which revealed that the 5’tRF^Glu(CTC)^ interactors form a highly interconnected network (**Figure S4.e**). Interestingly, RBPs and mRNA processing play a crucial role in β-cell homeostasis^18, 19^. Moreover, the RBP Msi2, detected as 5’tRF^Glu(CTC)^ interactor, was shown to be overexpressed in immature β-cells and to be induced in islets in conditions associated with T2D, such as ER stress and lipotoxicity, controlling mRNA translation^20^.

**Figure 5.**
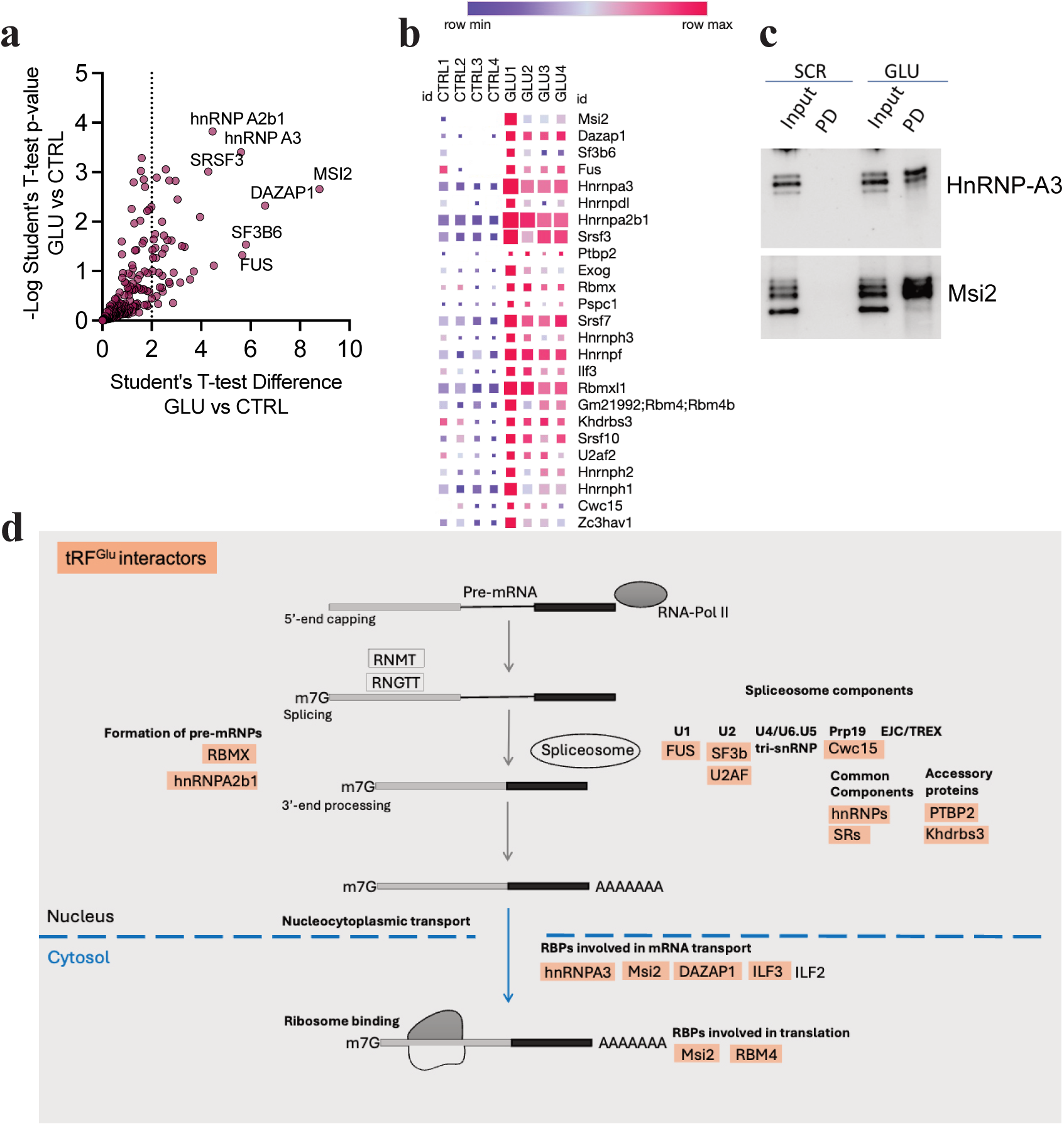
Identification of 5’tRF^Glu(CTC)^ interactors: MIN6 cells transfected with a biotinylated 5’tRF^Glu(CTC)^ mimic (GLU) or a scrambled control (CTRL) were used for pull-down and mass spectromety. (**a**) Volcano plot of the proteins enriched in the GLU pull-down versus CTRL. (**b**) Heatmap of the 25 proteins selected as possible interactors (enrichment FC>2 GLU mimic vs CTRL, unique peptides>2, p<0.05), the square size indicates the IBAQ values (0 to 14). (**c**) Western blot of hnRNP-A3 and MSI2 validating the mass spectrometry data. (**d**) The scheme represents the involvement of 5’tRF^Glu(CTC)^ interactors in the different steps of mRNA processing pathway.

Because of the identified interactions with RBPs, we further investigated the consequences of 5’tRF^Glu(CTC)^ modulation in transcriptional and post-transcriptional gene regulation. To this purpose, we conducted transcriptomic and proteomic analysis in mouse islet cells upon transfection with either Ctr or αGlu ASO (**Figure 6.a**).

**Figure 6.**
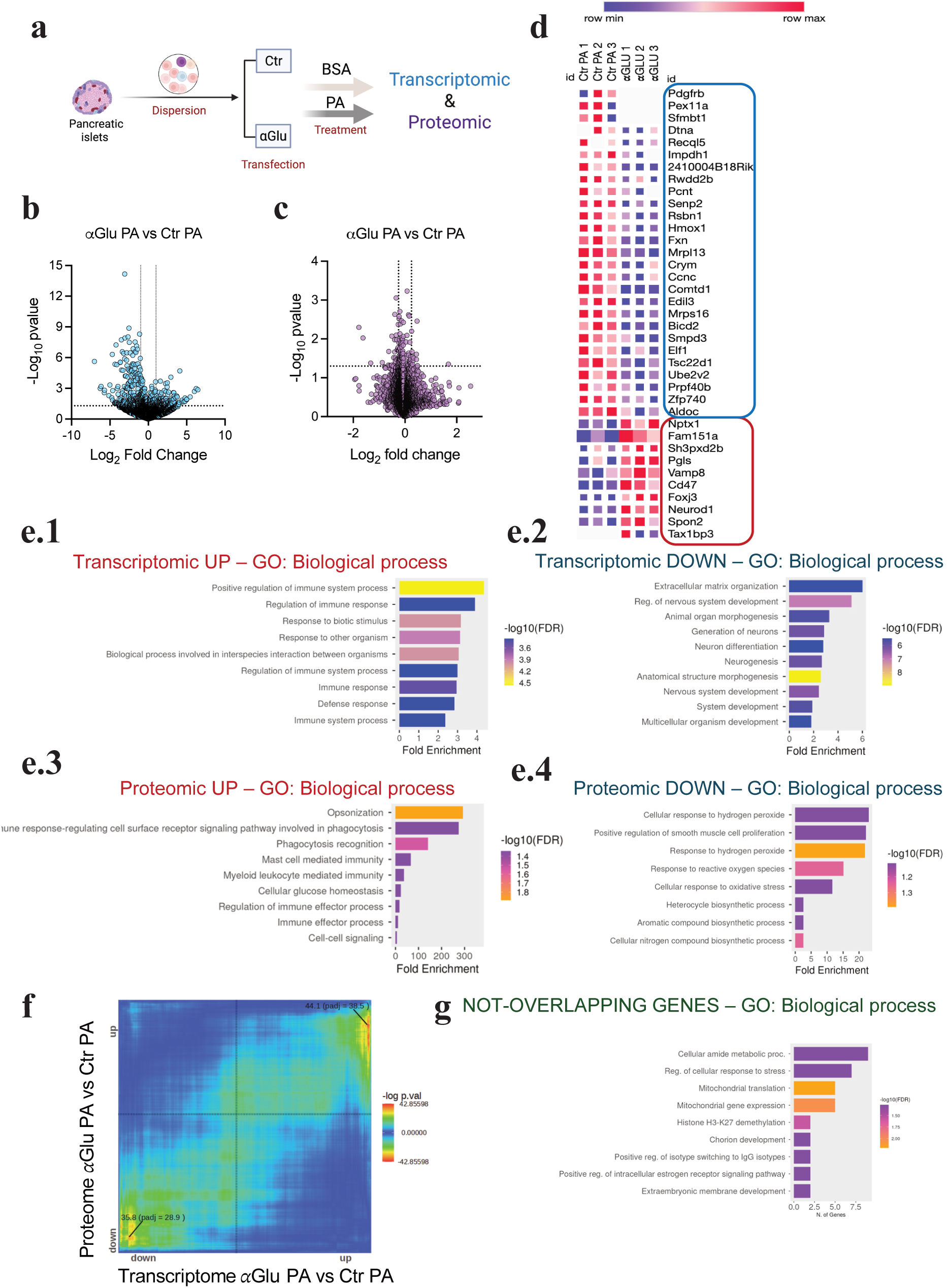
Analysis of gene regulation mechanisms affected by 5’tRF^Glu(CTC^) inhibition: Primary mouse islets cells were transfected with antisense oligonucleotide (ASO) of control (Ctr) or inhibiting 5’tRF^Glu(CTC)^ (*⍺*-Glu) and then treated with either BSA or palmitate (PA), gene and protein expression were assessed by transcriptomic and proteomic analysis (**a**). Volcano plots showing differential expressed genes (**b**) and proteins (**c**) in dispersed mouse islets due to 5’tRF^Glu(CTC)^ inhibition in palmitate treatment condition (**α**Glu PA vs Ctr PA). The proteins significantly modulated (p<0.05) by the tRF inhibition in palmitic condition are shown in the heatmap (**d**). (**e1-4**) Overrepresented GO biological process terms in up- and down-regulated genes (**e1-2**) and proteins (**e3-4**). Rank-rank hypergeometric overlap of the modulation in proteome and transcriptome of αGlu transfected cells under palmitate treatment was performed with RedRibbon (**f**). Functional enrichment of not-overlapping genes from rank-rank hypergeometric overlap analysis (**g**).

mRNA sequencing identified 1534 genes modulated in mouse islets in response to palmitate treatment (Ctr PA vs Ctr BSA, FC>2, p<0.05) (**Figure S5.a**), with a downregulation of genes involved in haematopoiesis and cell adhesion, and an upregulation of fatty acid metabolism components (**Figure S5.c1-4**). Upon 5’tRF^Glu(CTC)^ inhibition, we detected 434 differentially expressed genes (FC>2, p<0.05) in BSA (**Figure S5.b**) and 400 in palmitate-treated cells (PA) compared to controls (**Figure 6.b**). Interestingly, in both BSA and PA conditions, inhibition of 5’tRF^Glu(CTC)^ led to an upregulation of genes involved in monocyte attraction and in immune responses (**Figure 6.e1 and S5.d1**). Functional analysis of the downregulated genes in BSA condition revealed a significant enrichment of pathways related to antigen presentation and adaptive immune responses (**Figure S5.d2**). In PA condition, we observed a downregulation of genes involved in enteric nervous system development and extracellular matrix organization (**Figure 6.e2).** Although these genes are not directly linked to β-cell homeostasis, several of the modulated transcripts (e.g., *Gdnf, Ngfr, S100b, Hand2, Foxd3, Mrgprf, Cadm3, Gap43, Gpm6b, Cd109, Mmp17, Chst3, Gfap*) are associated with neurotrophic signaling and cell interaction pathways, indicating a potential role in the modulation of islet microenvironment.

Because of the apparent link between 5’tRF^Glu(CTC)^ and mRNA processing and translation, we performed a proteomic characterization of mouse islet cells upon PA exposure and 5’tRF^Glu(CTC)^ inhibition. Proteomic analysis identified 8359 proteins, 218 of which were modulated by PA treatment (Ctr PA vs Ctr BSA, FC>1.25, p<0.05) (**Figure S6.a**). In accordance with the transcriptomic data, we observed a downregulation of proteins involved in extracellular matrix composition and metabolic responses to glucose (**Figure S6.c1**) and an upregulation of proteins implicated in transport to peroxisome and fatty acid degradation (**Figure S6.c2**). We identified 82 differentially expressed proteins in α*-*Glu versus Ctr ASO in basal conditions (α*-*Glu BSA vs Ctr BSA, FC>1.25, p<0.05) (**Figure S6.b**) and 26 in PA exposure condition (α*-*Glu PA vs Ctr PA FC>1.25, p<0.05) (**Figure 6.c-d**). In agreement with the transcriptomic analysis, tRF inhibition led to the upregulation of proteins related to phagocytosis and innate immunity, in both BSA and PA conditions (**Figure 6.e3 and S6.d1**). The tRF blockade in basal conditions led to the downregulation of pathways linked to adaptive immune responses (**Figure S6.d2)**. Upon PA treatment, tRF blockade resulted in the downregulation of proteins involved in oxidative stress responses (**Figure 6.e4**). Notably, we found that some proteins modulated by PA exposure were maintained at their physiological levels in 5’tRF^Glu(CTC)^-depleted islet cells (**Figure S6.e1-3**). For instance, the drastic drop caused by PA treatment of Tax1bp3, a repressor of Wnt/β-catenin pathway, is completely prevented in cells lacking 5’tRF^Glu(CTC)^. Similarly, the downregulation of NeuroD1 and VAMP8, important for β-cell differentiation^21^ and insulin secretion^22^, respectively, occurring in PA-treated cells is no longer observed upon blockade of 5’tRF^Glu(CTC)^.

All together our functional investigations suggest that 5’tRF^Glu(CTC)^ contributes not only to endogenous β-cell processes, by regulating the levels of proteins involved in oxidative stress and insulin secretion, but also to islet dynamics, by controlling the expression of genes and proteins involved in innate immunity activation.

We then used a rank–rank hypergeometric overlap-based method to investigate the link between proteomic and transcriptomic regulation^23^. The proteomic and transcriptomic modulations induced by PA treatment showed a high overlap (**Figure S6.f**), as it has been previously shown in β-cell lines. On the other hand, the effect of 5’tRF^Glu(CTC)^ inhibition on protein and gene expression in PA condition was only partially overlapping (**Figure 6.f**). Interestingly, we found a group of genes with opposite modulation at the level of mRNA or protein expression (**Table S4**). While the mRNAs were upregulated or not modulated, the proteins encoded by these genes were downregulated by the 5’tRF^Glu(CTC)^ inhibition. These proteins are part of mitochondrial translation and stress responses (GO: biological processes) (**Figure 6.g**). These results suggest that tRFs may act as transcriptional but also post-transcriptional regulators.

### 5’tRF^Glu(CTC)^ modulates macrophage gene expression and polarization

As the blockade of 5’tRF^Glu(CTC)^ in mouse islet cells leads to the upregulation of genes and proteins involved in innate immune activation, we decided to further characterize the role of 5’tRF^Glu(CTC)^ in Mφs polarization. We first assessed the levels of the tRFs in bone marrow derived Mφs during *in vitro* polarization into pro- and anti-inflammatory phenotypes (**Figure S7.a1-5**). We found that 5’tRF^Glu(CTC)^ levels peak 24 hours after the induction of anti-inflammatory phenotype by IL-4 and IL-13 and progressively decline thereafter during pro-inflammatory polarization induced by LPS and IFN-γ (**Figure 7.a**).

**Figure 7.**
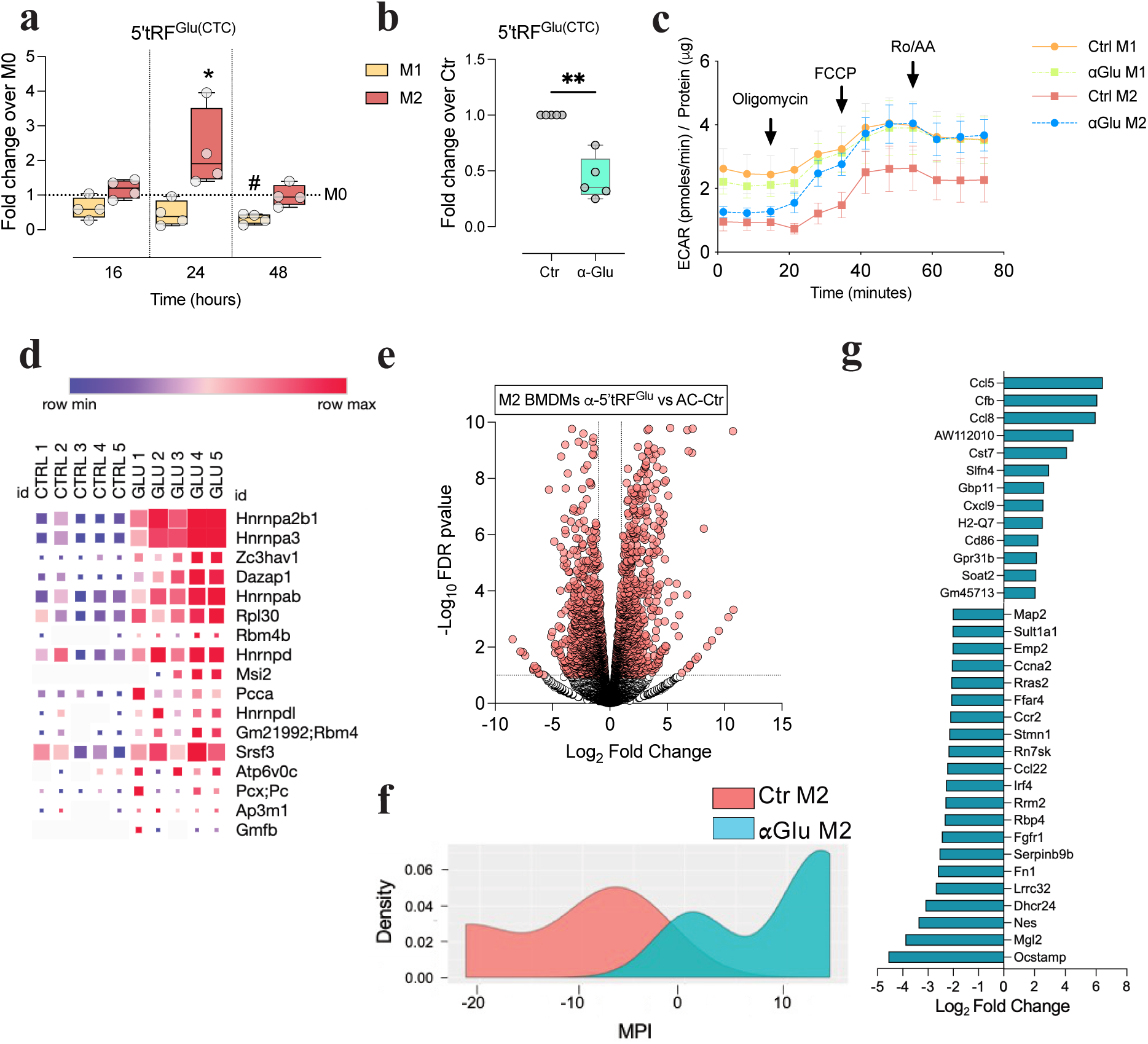
5’tRF^Glu(CTC)^ is necessary for M2 anti-inflammatory macrophage polarization: 5’tRF^Glu(CTC)^ levels were assessed by qPCR at the indicated time points of BMDM polarization (**a**). Transfection of naïve M0 macrophages with antisense oligonucleotide (ASO) targeting 5’tRF^Glu(CTC)^ (**α**Glu) or antisense of control (Ctr) was performed prior BMDM polarization (**b**). The extracellular acidification rate of M1 or M2 macrophages, transfected with either **α**Glu or Ctr antisense was assessed with Seahorse analyser during a mito-stress assay (**c**). Heatmap show the proteins identified as 5’tRF^Glu(CTC)^ interactors in M2 macrophages by mass spectrometry (enrichment FC>2 GLU mimic vs CTRL, unique peptides>2, p<0.05), the square size indicates the IBAQ values (0 to 12) (**d**). (**e-g**) Transcriptomic profiling of M2 macrophages upon 5’tRF^Glu(CTC)^ inhibition: volcano plot shows the differential expression of genes compared to M2 transfected with Ctr ASO (**e**). MacSpectrum tool was used to evaluate the macrophage polarization state based on their transcriptomic profile (**f**) and to identified the key genes involved in the polarization shift towards a pro-inflammatory phenotype observed in M2 **α**Glu compared to Ctr (**g**). *p<0.05by One-way ANOVA with Sidak correction for multiple comparisons; **p<0.01 by Student –t-test.

We next investigated whether 5’tRF^Glu(CTC)^ contributes to the establishment of an anti-inflammatory Mφ phenotype. To this end, we inhibited the expression of this fragment in M0 Mφs using an antisense oligonucleotide (α-Glu ASO) prior to polarization (**Figure 7.b**). To evaluate Mφ metabolic activity, we measured oxygen consumption rates (OCR) and extracellular acidification rates (ECAR) using the Seahorse Mito Stress Test (Agilent). OCR serves as a readout of mitochondrial respiration, which is typically elevated in anti-inflammatory Mφs, whereas ECAR provides an indirect measure of glycolysis, the dominant energy pathway in pro-inflammatory Mφs^24^. We found that blockade of 5’tRF^Glu(CTC)^ lead to higher ECAR in anti-inflammatory Mφs (**Figure 7.c**). At the same time, the OCR measurements showed an increased response to FCCP (**Figure S7.b**), indicating higher maximal respiratory capacity. This may be due to metabolic reprogramming^24^. Using a biotinylated mimic of 5’tRF^Glu(CTC)^, we pulled down the proteins interacting with the fragment in anti-inflammatory macrophages (**Figure 7.d and Figure S7.c-d**). We found 17 proteins enriched in the pull-down performed with the tRF mimic (GLU), 8 of which common to those pull-down in β-cells. As in β-cells, 5’tRF^Glu(CTC)^ interactors were mainly RBPs involved in mRNA splicing, processing and translation (**Figure S7.e1-2**). Therefore, we investigated whether 5’tRF^Glu(CTC)^ modulates gene expression in Mφs. mRNA sequencing of anti-inflammatory Mφs revealed that 5’tRF^Glu(CTC)^ inhibition causes major transcriptional changes (**Figure 7.e**) with the downregulation of genes involved in cell division and lipid metabolism and an upregulation of genes related to inflammatory pathways (**Figure S7.f1-2**). Using MacSpectrum, a tool that predicts Mφ phenotype based on transcriptomic data^25^ we found that 5’tRF^Glu(CTC)^ inhibition triggers a shift towards higher and more pro-inflammatory Mφ polarization index (MPI) (**Figure 7.f**) because of changes in the expression of several key regulatory genes (**Figure 7.g**). Our observations point to a central role of tRFs in determining Mφ activation via regulation of gene expression.

## DISCUSSION

In this study, we investigated the role of tRF modulation in pancreatic islets during the development of T2D. Our findings demonstrate that tRFs are dynamically modulated *in vivo* in islets from an obese and diabetic mouse model as well as from individuals with IGT and T2D; we showed that 5’tRF biogenesis is induced *in vitro* by free fatty acid exposure and identified 5’tRF^Glu(CTC)^ as a key regulator of islet gene expression, stress responses and macrophage activation.

### Modulation of tRF cleavage in islet homeostasis and disease

Recent studies have highlighted the role of tRFs as critical regulatory molecules in islet biology. tRNA cleavage was initially implicated in early-onset diabetes in patients carrying mutations in the tRNA-modifying enzyme TRMT10A^10^. Our previous work showed that tRFs are physiologically expressed in rodent islets and are modulated during postnatal β-cell maturation^12^ and in response to nutritional changes^13^. In conditions such as obesity and IGT, exposure to elevated glucose and circulating free fatty acids triggers sustained cellular stress, contributing to β-cell dysfunction and apoptosis. This stress is accompanied by the expansion and activation of islet macrophages, which monitor and modulate β-cell fitness.

Despite these insights, the functional relevance of tRFs in islet homeostasis, their modulation in human T2D and their role in islet cell crosstalk remained largely unexplored. Using small RNA sequencing, we found an upregulation 5’tRFs derived from tRNA^Gly(GCC)^ and tRNA^Glu(CTC)^ in islets from T2D patients and in both iMACs and β-cells from db/db mice. Notably, 5’tRF^Glu(CTC)^ was also upregulated in islets from individuals with IGT, with levels inversely correlating with insulin secretion and glucose sensitivity. These findings suggest that early induction of this tRF may contribute to diabetes onset. Interestingly, 5’tRF^Glu(CTC)^ is abundant in neonatal rat islets but downregulated during maturation^12^, suggesting that low levels of this fragment may be required to sustain mature β-cell function. Supporting this, downregulation of 5’tRF^Glu(CTC)^ in islet cells preserved residual glucose-induced insulin secretion following exposure to saturated fatty acids. We further demonstrated that this induction observed *in vivo* can be mimicked *ex vivo* by exposing islet cells to saturated fatty acids, especially palmitate.

The generation of tRFs under physiological conditions and their induction under stress is driven by differential expression of tRNA genes^26^, or of tRNA-cleaving enzymes^14, 15, 27^. We found that palmitate does not alter the level or the aminoacylation of tRNA^Glu(CTC)^ but is likely to promote its cleavage by increasing the expression of the endoribonuclease Angiogenin which is also upregulated in FACS sorted β-cells from db/db mice. However, the induction of 5’tRF^Gly(GCC)^ was not prevented by Angiogenin knockdown, suggesting either incomplete silencing, activation via post-translational modifications, such as phosphorylation and translocation^28, 29^, or the involvement of alternative RNAses.

### tRF function in islet stress responses and iMAC activation

T2D typically arises from a pre-diabetic state characterized by impaired glucose tolerance (IGT) and mild hyperglycemia, with obesity being a key risk factor. Chronic nutrient excess initially triggers compensatory β-cell mass expansion and the activation of stress responses but eventually leads to β-cell failure. We found that inhibition of tRF^Glu(CTC)^ in dispersed islet cells prevents palmitate-induced apoptosis and restores the residual capacity of β-cells to release insulin in response to glucose, suggesting a direct role in stress-induced β-cell dysfunction.

Obesity and T2D are also chharacterized by chronic low-grade inflammation. This has been extensively studied in adipose tissue, where macrophages recruited from the circulation adopt a pro-inflammatory phenotype and release cytokines and extracellular vesicles that can affect islet function^30, 31^. In physiological conditions, islet macrophages exhibit an activation state characterized by the production of pro-inflammatory cytokines^16^ but also by the expression of homeostatic genes^3^. In obesity, these cells undergo transcriptional reprogramming toward a more homeostatic phenotype marked by decreased IL-1β and increased expression of anti-inflammatory genes^3, 6^.

Using a co-culture system, we modeled the activation dynamics of iMACs observed in mouse models of T2D. Bone marrow-derived macrophages co-cultured with MIN6 β-cells acquired an iMAC-like phenotype and, when exposed to palmitate, transitioned to an obesity-associated activation state. This indicates that β-cells influence macrophage polarization and that macrophage precursors can acquire iMAC characteristics under appropriate stimuli. While embryonic origin may be required for the acquisition of a fully specialized phenotype, precursor-derived macrophages may contribute to the maintenance of the iMAC pool. Our co-culture system enables studies on iMAC-like cell functions and on their crosstalk with β-cells. Indeed, we demonstrated that blockade of tRF^Glu(CTC)^ in macrophages prevents palmitate-dependent activation and protects β-cells in co-culture. These findings were corroborated by observations in polarized bone marrow-derived macrophages, where blockade of this tRF in anti-inflammatory macrophages led to a shift to a hypermetabolic, pro-inflammatory phenotype.

### Molecular mechanisms of tRF action

Gene regulation under stress involves the rapid and coordinated activity of multiple regulatory molecules. Small non-coding RNAs are rapidly modulated by environmental cues and their interaction with RBPs can reshape both transcriptomic and proteomic landscapes. We identified a network of RBPs interacting with tRF^Glu(CTC)^, many of which are involved in mRNA processing and known to play roles in islet homeostasis and stress adaptation^19, 20^. Consistent with these interactions, blockade of tRF^Glu(CTC)^ in islet cells altered gene and protein expression under both basal and lipotoxic conditions. Upon palmitate exposure, inhibition of tRF^Glu(CTC)^ led to upregulation immune activation pathways, suggesting that this tRF normally acts to mitigate inflammatory responses under diabetogenic conditions by promoting a more homeostatic iMAC state. Additionally, depletion of tRF^Glu(CTC)^ downregulated genes involved in extracellular matrix remodeling and neurogenesis, processes likely to be important in reshaping the islet microenvironment during lipotoxicity^32, 33^.

Proteomic analysis revealed that tRF^Glu(CTC)^ inhibition reduced the expression of proteins involved in oxidative stress responses, suggesting that these effects may underlie the protective effects observed in β-cells, since induction of oxidative stress is considered a major contributor of islet cell demise during lipotoxicity^17^.

Although some pathways were consistently modulated at both mRNA and protein levels, others showed discordant patterns, likely reflecting post-transcriptional control. Rank-rank hypergeometric overlap analysis confirmed that palmitate induced robust and concordant transcriptomic and proteomic changes^34^, while the effects tRF inhibition were more selective, particularly affecting genes related to stress responses and mitochondrial translation.

### Conclusions

Our study uncovers a novel regulatory framework in which tRFs — particularly 5’tRF^Glu(CTC)^ — modulate islet responses to diabetogenic stress. In pre-diabetic and early T2D diabetes states, tRFs may act as integrators of environmental signals, coordinating endocrine and immune responses and contributing to islet adaptation or dysfunction.

### Limitations of the study and perspectives

Our results suggest that tRFs respond to metabolic cues and orchestrate islet cell function and intercellular communication. While we focused on 5’tRF^Glu(CTC)^, other fragments such as 5’tRF^Gly(GCC)^ also showed modulation and warrant further study. Future investigations should delineate the specific role of these fragments in islet biology and macrophage activation. In addition, the precise mechanisms by which tRFs influence RBP activity and mRNA processing remain to be elucidated in detail.

Moreover, while our co-culture system captures some essential aspects of iMAC-β-cell interactions, more physiologically relevant 3D models are needed to explore the complex islet microenvironment dynamics. Recent studies using embryonic stem cell-derived organoids have highlighted the importance of iMACs in β-cell maturation^35^. Extending tRF profiling and manipulation to such models could yield critical insights into their roles in islet development and function.

Finally, *in vivo* delivery of ASOs to target and block tRFs in specific islet cells represents a major and challenging perspective. Targeted delivery to β-cells using glucagon-like peptide-1 receptor (GLP1R) ligand carriers has shown encouraging results^36, 37^ and ongoing work is exploring the use of scavenger receptor ligands to reach tissue-resident macrophages^38, 39^. Notably, ASO uptake by activated macrophages and subsequent delivery to parenchymal cells has been described^40, 41^, highlighting the importance of studying macrophage-cell interactions in the context of tRF-targeted therapies.

## Experimental models and subject details

### Mice

All animal procedures and protocols were approved by the Swiss Research Councils and Veterinary Offices under the animal authorization number VD2495×4.

Db/db mice (genetic background BKS.Cg-*Dock7^m^*+/+*Lepr^db^*J) and wild type controls (genetic background BKS.Cg-*Dock7^m^*+/+) of 8 weeks of age were obtained from Charles River Laboratories (Les Oncins, France). C57BL/6NRj mice (aged 12-14 weeks) were obtained from Janvier Laboratories (Le Genest-Saint-Isle, France) or derived from our colony. Mice were housed on a 12-h light/dark cycle under standard conditions with *ad-libitum* chow diet.

### Living organ donors

Living donors were metabolically profiled before undergoing surgery, as previously described^42, 43^, and subjected to a 75 g OGTT and HbA1c testing to exclude diabetes, according to the American Diabetes Association criteria^44^. None of the participants enrolled had a family history of diabetes.

A standard 75 g OGTT was performed after a 12 h overnight fast with measurements of glucose, insulin and C-peptide at 0, 30, 60, 90 and 120 min after the glucose load. Based on the OGTT, we classified the population as normal glucose tolerant (NGT n=12), impaired glucose tolerant (IGT n=13) and having T2D (T2D n=11), according to the American Diabetes Association criteria^44^. During OGTT, insulin secretion was derived from C-peptide levels by deconvolution. Beta cell glucose sensitivity, i.e. the slope of the relationship between insulin secretion and glucose concentration, was estimated from the OGTT by modelling, as previously described^45, 46^.

## Methods

### Cell lines

MIN6B1 cells, a murine insulin-secreting cell line (Lilla et al., 2003), were cultured at a density of 1.5 × 10^5^ cells/cm^2^ in DMEM-GlutaMAX medium (Invitrogen) containing 25 mM glucose and 4 mM L-glutamine, and supplemented with 15% fetal calf serum, 70 mM b-mercaptoethanol, 50 mg/mL streptomycin and 50 IU/mL penicillin. All cell lines were cultured at 37°C in a humidified atmosphere (5% CO2, 95% air) and tested negative for mycoplasma contamination.

### Mouse islet isolation and cell sorting

Mouse pancreases were digested with collagenase (Sigma), and islets were isolated by Histopaque (Sigma) density gradient and handpicking^47^. For the indicated experiments, mouse islets were dissociated into single cells by incubation in Ca^2+^/Mg^2+^ free phosphate-buffered saline, 3 mM EGTA, and 0.002% trypsin (ThermoFisher) for 2-3 min at 37 °C. Isolated mouse islets and dissociated cells were cultured in RPMI 1640 GlutaMAX medium (ThermoFisher) supplemented with 10% FCS, 100 U/mL penicillin and 100 µg/mL streptomycin, 1 mM sodium pyruvate (Sigma), and 10 mM Hepes (Sigma).

For fluorescence-Activated Cell Sorting (FACS), islets from two db/db mice and three wild type mice were pooled; 600-800 islets per preparation were dissociated. Fractions enriched of β-cells were obtained based on β-cell autofluorescence, as previously described^48^. Islet-resident macrophages (iMACs) were sorted based on the expression of CD45, Cd11b, F4/80 and Cd11c markers^6^. Dissociated islet cells were washed once with *FACS buffer* (0.1% BSA, 2mM EDTA, 11mM glucose in PBS) and incubated for 5 minutes with TruStain FcX™ (anti-mouse CD16/32) Antibody (BioLegend), at 4°C. Then cells were incubated for 30 min in the dark at 4°C with the following antibodies: 1:200 of FITC anti-CD45, 1:100 brilliant violet CD11b, 1:100 APC F4/80 and 1:100 PE CD11c (BioLegend). Cells were then washed twice with *FACS buffer* and sort by FCF-Aria-II (SORP). β-cell purity was assessed as previously described^49^. On average, β-cell fractions contained 99.1 ± 0.9% insulin-positive cells and 0.6 ± 0.6% glucagon-positive cells.

### Bone marrow derived macrophage differentiation

Bone marrow was isolated from tibia and femurs of 10-12 weeks old C57Bl/6N mice. Red blood cells were lysed in RBC lysis buffer containing 0.155 M NH4Cl, 10 mM KHCO3, 0.127 M EDTA. Bone marrow cells were differentiated for 7 days on not-tissue culture treated 100mm petri dishes in basal BMDM medium composed by DMEM (ThermoFisher 61965-059), 10mM HEPES, 10% FCS, 50 μg/mL streptomycin and 50 IU/mL penicillin, supplemented with 10ng/ml macrophage colony stimulating factor (M-CSF, Peprotech 315-02). After 7 days, bone marrow-derived macrophages (BMDMs) were further polarized into anti-inflammatory M2 macrophages by exposure to 10ng/ml IL-4 and IL-13 for 48 hours, and pro-inflammatory M1 macrophages by exposure to 100ng/ml lipopolysaccharide (LPS) and 50ng/ml IFN-γ for 48 hours. Differentiation and polarization were monitored by the expression of specific gene markers measured by qPCR.

### Pancreas samples and laser capture microdissection of islets from living donors

Pancreas samples were collected from living donors undergoing pylorus-preserving pancreatoduodenectomy recruited at the Digestive Surgery Unit and studied at the Centre for Endocrine and Metabolic Diseases Unit (Agostino Gemelli University Hospital, Rome, Italy).

Indications for surgery were periampullary tumors, pancreatic intraductal papillary tumors, mucinous cystic neoplasm of the pancreas, and nonfunctional pancreatic neuroendocrine tumors. Surgical pancreatic tissue specimens were snap frozen in liquid nitrogen and stored at −80°C embedded in Tissue-Tek OCT compound. The study protocol (ClinicalTrials.gov NCT02175459) was approved by the Ethical Committee Fondazione Policlinico Universitario Agostino Gemelli IRCCS – Università Cattolica del Sacro Cuore (P/656/CE2010 and 22573/14), and all participants provided written informed consent, followed by a comprehensive medical evaluation, as previously described^42, 43^.

Pancreatic human tissue samples from living donors were cryosectioned. Sections were fixed in 70% ethanol for 30 s, dehydrated in 100% ethanol for 1 min, in 100% ethanol for 1 min, in xylene for 5 min and finally air-dried for 5 min. Laser capture microdissection (LCM) was performed using an Arcturus XT Laser-Capture Microdissection system (Arcturus Engineering, Mountain View, CA, USA) by melting thermoplastic films mounted on transparent LCM caps (cat. LCM0214 - ThermoFisher Scientific, Waltham, MA, USA) on specific islet areas. Human pancreatic islets were subsequently visualized through islet intrinsic autofluorescence for LCM procedure. Thermoplastic films containing microdissected cells were incubated with 10 µl of extraction buffer (cat. kit0204 - ThermoFisher Scientific, Waltham, MA, USA) for 30 min at 42 ◦ C and kept at −80°C until RNA extraction. Each microdissection was performed within 30 min from the staining procedure in a contamination-free, dehumidified environment maintaining the external temperature at 16°C to preserve RNA integrity. Overall, an endocrine area of microdissected islets of about 2×106 um2 for each subject was analyzed for the global molecular analysis.

### Human islet isolation from cadaveric donors

Human islets from brain-dead donors were obtained from the Centre d’Etude Européen pour le Diabéte in Strasbourg. The use of human tissue and all experimental protocols were approved by the French government (Directorate General for Research and Innovation of the Ministry, Bioethics Unit) registered under number DC-2019-3439. Informed consent was obtained from all subjects’ legal guardians. Upon arrival, human islets were dissociated and cells were cultured in CMRL medium (Invitrogen) supplemented with 10% fetal calf serum (Gibco), 10 mM HEPES pH 7.4, 100 mg/mL streptomycin and 100 IU/mL penicillin and 2 mmol/l glutamine. Detailed information about human islet donors is provided in Table S5.

### β-cell and macrophage co-culture

After differentiation in p100 dishes, BMDMs were detached with Accutase (Thermofisher) and plated in direct co-culture with MIN6 cells at the ratio of 1:10. After 24 hours co-cultures were treated either with 0.5 mM palmitate or with 0.75% BSA as control for additional 48 hours. Cells were either fixed with 4% PFA for immunofluorescence staining or used for insulin secretion assay. For gene expression analysis, the two cell populations were purified by FACS by staining macrophages with antibodies against F4/80 and CD45.

### Cell transfection and treatment

For tRF inhibition, MIN6 cells, primary mouse and human islets, BMDMs were transfected with 50 pmol custom miRCURY LNA miRNA inhibitor (Qiagen) and 2 μl of lipofectamine 2000 in 24 well plate. For Angiogenin silencing, MIN6 cells were transfected with 20 pmol of stealth siRNA (ThermoFischer) and 2 μl of lipofectamine 2000 in 24 well plate. After overnight incubation, medium was replaced with complete medium or with treatment medium. Oleic, stearic and palmitic acid were added at 0.5 mM concentration complexed with fatty-acid free BSA (Sigma - Aldrich). Control treatment was performed in 0.75% fatty-acid free BSA. Fatty acids treatments were performed in 5% FBS medium for MIN6 and BMDMs, and 1% FBS medium for primary cells.

### RNA purification

Total RNA was extracted from each LCM sample using PicoPure RNA isolation kit Arcturus (cat. kit0204 - ThermoFisher Scientific, Waltham, MA, USA) following manufacturer’s procedure. The characterization of RNA samples was performed with Agilent 2100 Bioanalyzer technology with RNA Pico chips (cat. 5067-1513 Agilent Technologies, Santa Clara, CA, USA): we considered as good quality samples those that showed a RIN higher or equal than 4.5 (scale: 1-10). Total RNA was extracted from mouse and human primary islet cells and macrophages, FACS-sorted cells and MIN6 cells using the miRNeasy micro kit (Qiagen). For sequencing applications, RNA with RIN higher than 7 was used.

### Real-time qPCR

Real-time PCR quantification of miRNAs and tRFs. was performed using the miRCURY LNA Universal RT microRNA PCR system. The sequences used for custom primer design are indicated in Table S6. For gene expression quantification, RNA was treated with DNase and reverse transcribed with a Moloney Murine Leukemia Virus reverse transcriptase and random primers. Quantitative PCR (qPCR) was performed using SsoAdvanced Universal SYBR Green Supermix. Primer sequences are listed in Table S6. Expression of miRNAs, tRFs and protein-coding genes was corrected for the level of miRNAs/ncRNAs and housekeeping genes that are unaffected by the experimental condition under study.

### Assessment of full length tRNA levels and aminoacylation

The tRNA levels and tRNA amino-acylation rates were assessed by a qPCR-based methodology based on a previously published protocols^50^. RNA was purified by phenol-chloroform extraction and precipitated with isopropanol. Samples were then resuspended in 10μl of cold tRNA resuspension buffer (10 mM sodium acetate buffer pH 4.5, 1 mM EDTA) and concentration was measured at nanodrop. From the same sample, 2 μg of RNA were incubated with 100 mM Sodium Acetate buffer in the presence of 10 mM NaIO4 (for “oxidized reaction”) or NaCl (for “control”). In the “oxidized reaction”, amino-acylated tRNAs are protected, while uncharged tRNAs lose their 3’A residue. After 20 minutes at room temperature, reactions were quenched with 125 mM glucose. After adding a spike-in oligonucleotide, RNA was precipitated with 1 μl glycogen, 50 mM NaCl and 100% EtOH. In order to induce tRNA de-aminoacylation, RNA pellets were resuspended in 100μl of 50 mM Tris pH 9 and incubate at 37°C for 45 minutes and then quenched with 50 mM sodium acetate buffer (pH 4.5) and 100 mM NaCl. After overnight precipitation with 100% EtOH samples were cleaned up with Zymo kit, dissolved in 5μl RNAse-free water. 800 ng of RNA was used for adaptor ligation with T4 RNA Ligase2 KQ (NEB) and then for retro transcription with SuperScript^TM^ RT IV (Thermo Fisher). cDNA was amplified with specific primers to detect tRNA^Glu(CTC)^ and amino-acylation was assessed by the difference in Ct values between oxidized and control reaction samples.

### Small RNA sequencing

cDNA libraries were prepared with QIAseq miRNA kit (Qiagen) from RNA of FAC-sorted β-cells and iMACs of db/db and wild type mice and from LCM-islets of living donors. Single-end sequencing was performed on NovaSeq instrument (Illumina).

### tRF annotation and analysis

Following adapter sequences removal, identical reads longer than 16 nts were collapsed based on Unique Molecular Identifiers and aligned to the mouse genome (GRCm38.p6) or the human genome (hg38). Aligned reads were mapped to the mature tRNA sequences from the GtRNAdb database (http://gtrnadb.ucsc.edu/) and to the 22 mouse mitochondrial tRNA sequences from https://www.ncbi.nlm.nih.gov using bowtie (version 1; http://bowtie.cbcb.umd.edu). Bowtie parameters were set to output only perfect matches to tRNA sequences. Differentially expressed tRFs were identified using edgeR (version 4.2.2) in R packages (http://bioconductor.org). For each tRF, *p* values and false discovery rates (FDRs) were obtained based on the model of negative binomial distribution. Fold changes of tRF expression were also estimated within the edgeR package.

### Correlation analysis

Associations between the levels of tRFs detected by small RNA sequencing and clinical parameters of living donors were evaluated using linear models. tRFs normalized counts were converted in log2 scale after the addition of a pseudo count. To exclude influent values from the regression analysis, points with a cooks’ distance > 5 folds from the average cooks’ distance of the points in the regression model were removed. Linear models were then fitted for each tRF with each clinical parameter, implementing age, gender, and BMI as covariates. Regressions with a p-value associated to the coefficient of the clinical parameter <0.05 were considered as statistically significant. For representative purposes, the log2-transformed counts were corrected for the effect of the covariates for each regression. The correction was performed by estimating the effect of the covariates in each regression as the coefficient assigned to the covariate multiplied by the value of the covariate. Then, the effect of the covariates for each data point in the regression was summed up and then subtracted to the normalized log2-scaled counts.

### Pull-down and proteomics analysis

MIN6 cells were plated in 100 mm dishes and transfected with 20 nM biotinylated 5’tRF^Glu(CTC)^ mimic (GLU) or scramble control (CTRL) using 50 µl Lipofectamine 2000 (Thermo Fisher). After 48 h, cells were washed in cold PBS, crosslinked at 254 nm (200 mJ/cm²), and lysed in harsh buffer (150 mM NaCl, 50 mM Tris-HCl pH 7.5, 1% NP40, 0.5% sodium deoxycholate, 0.1% SDS). Lysates were clarified by centrifugation (10 min, 13,000 g), and incubated with M-280 streptavidin beads (Thermo Fisher). Inputs were stored. Proteins were processed using the SP3 method: eluted with 2% SDS/10 mM DTT/50 mM Tris pH 7.5, alkylated with 32 mM iodoacetamide, precipitated on Sera-Mag beads (ethanol 60%), and digested with trypsin. Peptides were desalted on SCX plates and analyzed by nanoLC-MS/MS (Vanquish Neo–Orbitrap Exploris 480, 130 min C18 gradient, DDA-HCD).

For M2 macrophages, bone marrow-derived M0 cells were polarized with IL-4/IL-13 for 48 h, lysed in harsh buffer, and incubated with GLU or CTRL mimics for 1 hour at RT before pulldown on streptavidin beads. Protein digestion followed the miST protocol: deoxycholate buffer (1% SDC, 100 mM Tris pH 8.6, 10 mM DTT), heat denaturation, Trypsin/LysC digestion, chloroacetamide alkylation, overnight digestion, SCX desalting. Peptides were separated on a 45 cm C18 column (140 min gradient) and analyzed on a Fusion Orbitrap (DDA-HCD).

Raw data were processed with MaxQuant 2.4.7.0 against the *Mus musculus* UniProt database (March 2023), with 1% FDR. iBAQ values were log2-transformed, filtered, and analyzed via Student’s t-test with Benjamini-Hochberg correction (FDR < 0.05). Complete protocols are provided in Supplementary Methods.

### mRNA sequencing

Libraries were prepared from total RNA with the Illumina Stranded mRNA Library Prep kit and sequenced with an AVITI or a NovaSeq6000 instrument using the SBS chemistry v4 (Illumina). After demultiplexing, reads annotation and differential expression analysis were performed using the QIAGEN RNA-seq Analysis Portal v 5.1.

### Proteomics

Cell lysates in RIPA buffer were processed using the SP3 protocol^51^. After dilution in SDS/DTT buffer and heat denaturation, proteins were alkylated and precipitated onto Sera-Mag Speedbeads with ethanol. Digestion was performed with trypsin, followed by desalting on SCX plates. Peptides were analyzed on a TIMS-TOF Pro mass spectrometer coupled to an EvoSep One LC system using a DIA-PASEF method with a 68 min gradient. Ion mobility and mass range selection followed established parameters^52^. DIA data were analyzed with Spectronaut 19.9 using the Pulsar engine and a *Mus musculus* UniProt reference proteome (February 2025). Peptides were quantified by XIC area, and protein groups inferred using the MaxLFQ algorithm. After log2 transformation and filtering, missing values were imputed, and differential expression was evaluated using Student’s t-test with Benjamini-Hochberg correction (FDR < 0.05). Detailed methods are available in Supplementary Methods.

### Bioinformatic tools

Gene and protein lists were analysed for pathway enrichment using ShinyGO 0.82 tool^53^. KEGG and GO enrichments with FDRpvalue<0.1 were considered significant. Rank-rank hypergeometric overlap was performed using the R-package RedRibbon1.3.1^23^. Transcriptomic data of M2 macrophages were run on MacSpectrum web tool^25^.

### Western blot and immunofluorescence

Proteins from cellular lysates were migrated on 10% SDS-PAGE gels. Following electrophoresis, proteins were transferred onto PVDF membranes and blocked at room temperature in Tris-buffered saline/0.1% Tween20 containing 5% of BSA. The membranes were then incubated overnight at 4°C with in 5% BSA. The bands were detected by chemiluminescence (Pierce) after incubation with a horseradish peroxidase-conjugated secondary antibody (Bio-Rad).

For immunofluorescence, cells were plated on chamber slide wells (Ibidi) and fixed with 4% paraformaldehyde. Cells were permeabilized with 90% methanol, blocked in 1X TBS buffer with 5% bovine serum albumin (BSA) and 0.1% Triton. Slides were then incubated with primary antibodies against mouse insulin (Proteintech) and cleaved caspase 3 (Cell signalling) and signal was detected with fluorescent secondary antibodies (ThermoFisher).

### Insulin secretion

MIN6B1 alone or in co-culture with iMAC-like macrophages were pre-incubated for 50 min at 37°C in Krebs-Ringer bicarbonate buffer (KRBH) containing 25 mM HEPES, pH 7.4, 0.1 % BSA (Sigma-Aldrich) and 2 mM glucose. The cells were then incubated for 50 min in KRBH containing 0.5% BSA and 2 mM glucose and the media were collected (basal insulin secretion). The cells were incubated for 50 min in KRBH containing 0.5% BSA and 20 mM glucose and the media were collected (stimulated insulin secretion). Total cellular insulin contents were recovered in acid-ethanol (0.2 mM HCl in 75% ethanol). Insulin levels were measured using an insulin enzyme immunoassay kit (Mercodia). For normalization with protein content, cells were lysed in RIPA buffer and proteins were quantified by bradford assay (Thermo Fischer). All experiments were performed in quadruplicates.

## Acknowledgments

This research was supported by the European Union’s Horizon 2020 research and innovation program under the Marie Skłodowska-Curie grant agreement No. 101029421 – MATREX to C.C.; the Transition Grant from the faculty of Biology and Medicine, University of Lausanne to C.C.; Swiss National Science Foundation - grant 310030_219252 to R.R.. We are grateful to Prof. Ping-Chih Ho (University of Lausanne) for technical and strategical advice on the analysis of macrophage immunometabolism. We thank Dr. Cédric Gobet, Dr. Lina Worpenberg and Prof. Felix Naef (École Polytechnique Fédérale de Lausanne) for sharing their knowledge and protocols for the analysis of tRNA levels and aminoacylation. We are grateful to Dr. Giulia Gliozzo, Dr. Lorenzo Carciero, Dr. Laura Soldovieri, Dr. Michela Brunetti and Dr. Emanuele Gentile (Catholic University of Sacred Heart, Rome, Italy) for technical assistance in human pancreas samples collection and in-vivo clinical metabolic evaluation. We are obliged to the numerous colleagues who contributed with insightful discussion and shared knowledge on various aspects of this study, notably Dr. Bilal Bayazit, Dr. Cécile Jacovetti (University of Lausanne) and Prof. Mariana Igoillo-Esteve (University of Brussels).

## Author contributions

Conceptualization, C.C., R.R.; Data curation, C.C., E.A, S.A., T.M., A.M.; Formal analysis, C.C., E.A., S.A., E.M., A.M.; Funding acquisition, C.C., R.R., Investigation C.C., R.K, V.M., C.G., E.A., D.A, G.D.G., G.C., A.G, F.A, F.B,; Methodology, C.C., R.K., V.M., E.A., G.D.G., G.C., A.M, S.A.; Project administration, C.C.,R.R.; Resources, C.C., R.R., T.M., G.Q., S.A., A.G. G.S., F.D.; Supervision, C.C., R.R., S.A, S.C., A.G., T.M., F.D. S.G.,; Validation, C.C., R.R., Visualization, C.C., R.R., T.M.; Writing – original draft, C.C.; Writing – review & editing, C.C., R.R., C.G., A.G., F.A., F.B, S.A, S.C, T.M., F.D., G.S.

## Supplementary Figures and Legends

**Figure S1:**
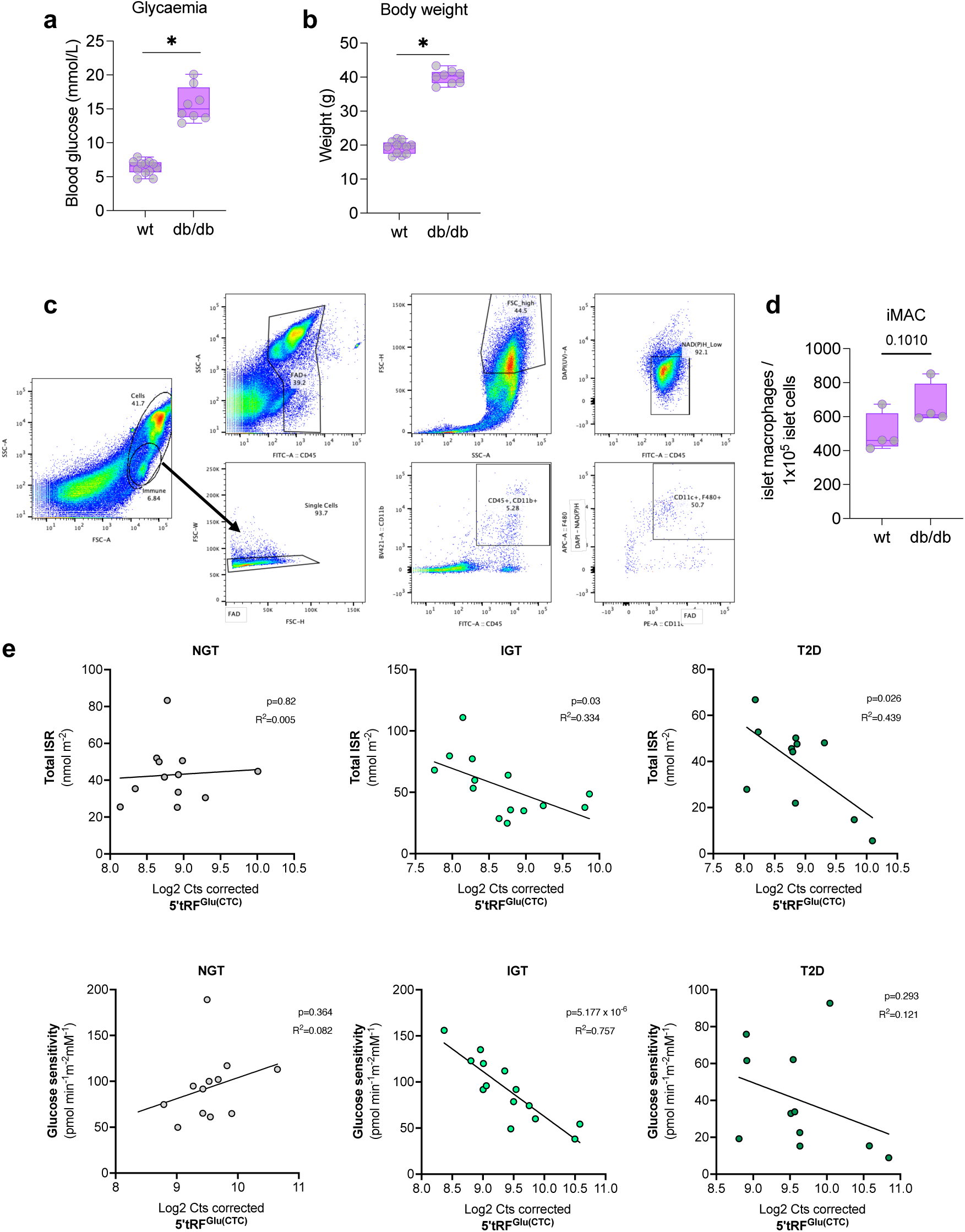
Blood glucose concentration (**a**) and body weight (**b**) were measured in 8 weeks old db/db mice and lean wild type (wt) controls. Gating strategy used for parallel FACS sorting of islet macrophages (iMACs) and *β*-cells (**c**). The numbers of iMACs from wt and db/db islets were plotted in (**d**). The levels of 5’tRF^Glu(CTC)^ detected in small RNA sequencing from patient subgroups were correlated with clinical parameters (**e**). *p<0.05 by Student t-test.

**Figure S2:**
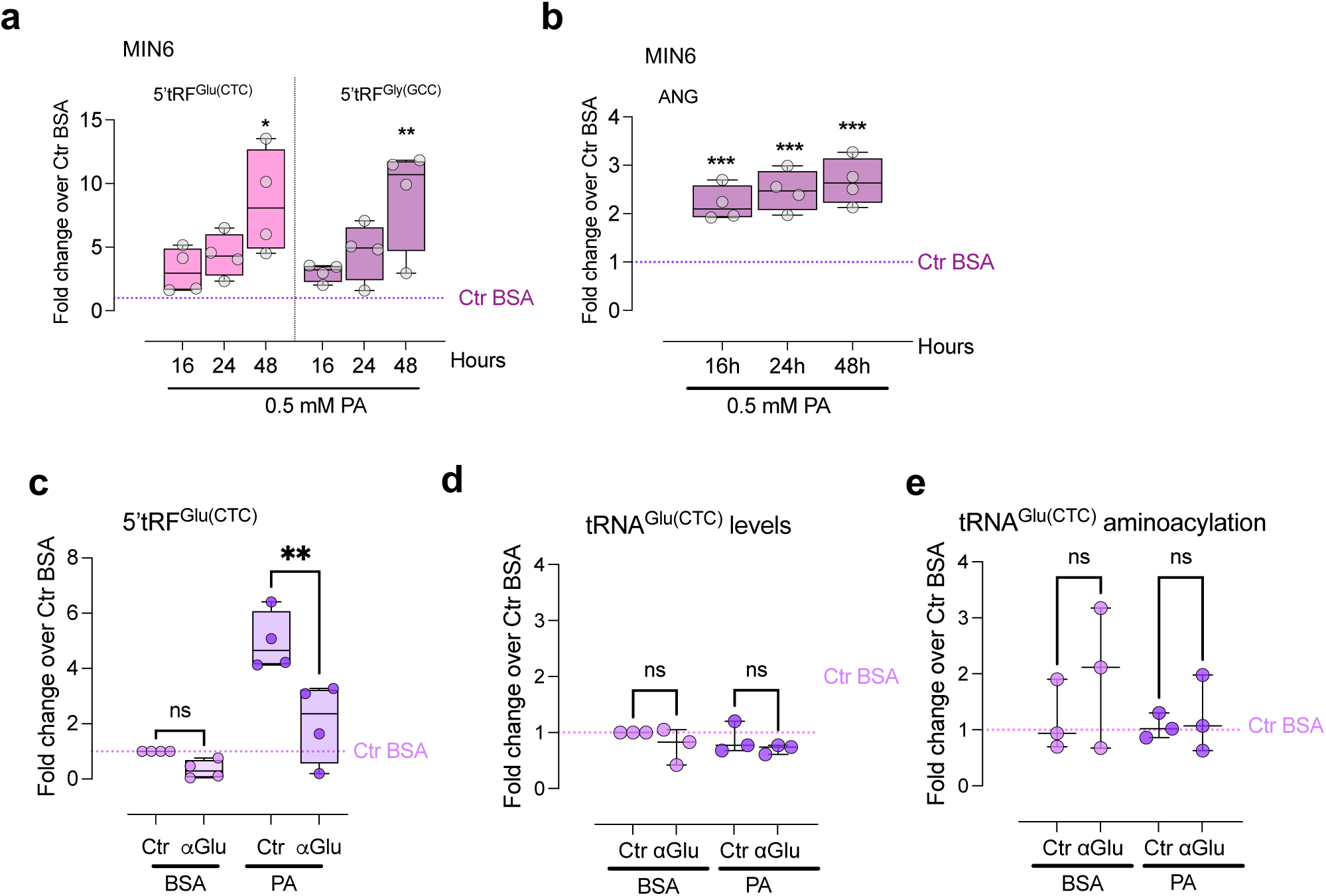
The expression of 5’tRFs (**a**) and Angiogenin (*ANG*) mRNA (**b**) was assessed by qPCR in MIN6 *β*-cells in different time points of palmitate exposure. Following **α**Glu ASO transfection in MIN6 cells, the levels of the 5’tRF^Glu(CTC)^ (**c**) and of the full length tRNA^Glu(CTC)^ (**d**) were measured. The aminoacylation rates of tRNA^Glu(CTC)^ were also assessed in the different conditions (**e**). *p<0.05; **p<0,01; ***p<0.001 by One-way ANOVA with Sidak correction for multiple comparisons.

**Figure S3:**
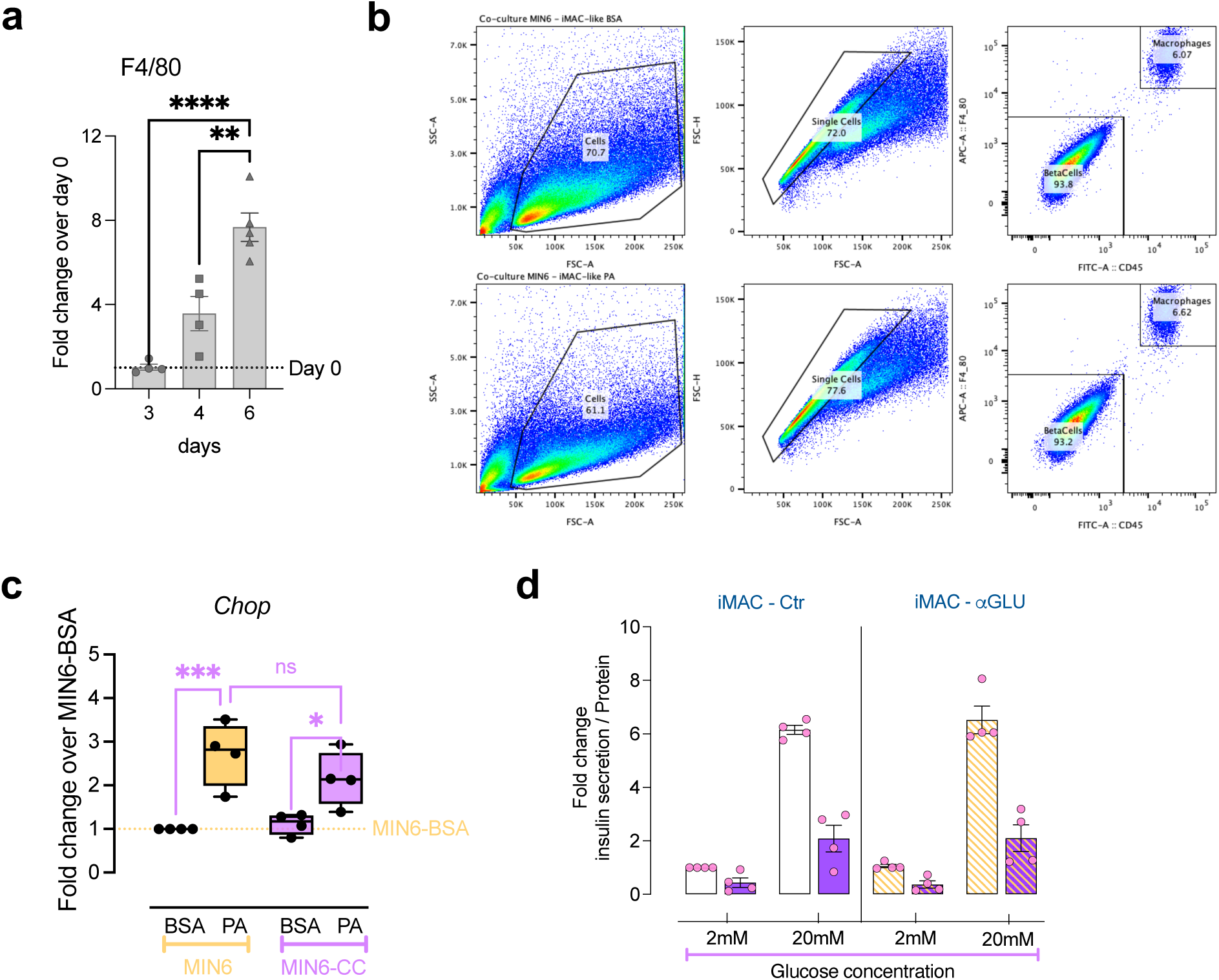
Bone marrow derived macrophages differentiation was assessed by the expression of *f4/80* (**a**). FACS gating strategy was used to separate macrophages and beta cells at the end of co-culture experiments (**b**). Expression of *Chop* (**c**) and insulin secretion normalized by protein content (**d**) in Min6-*β*cells cultured alone (yellow bars) or after co-culture with macrophages (pink bars). *p<0.05, ***p<0.001 by One-way ANOVA with Sidak correction for multiple comparisons.

**Figure S4:**
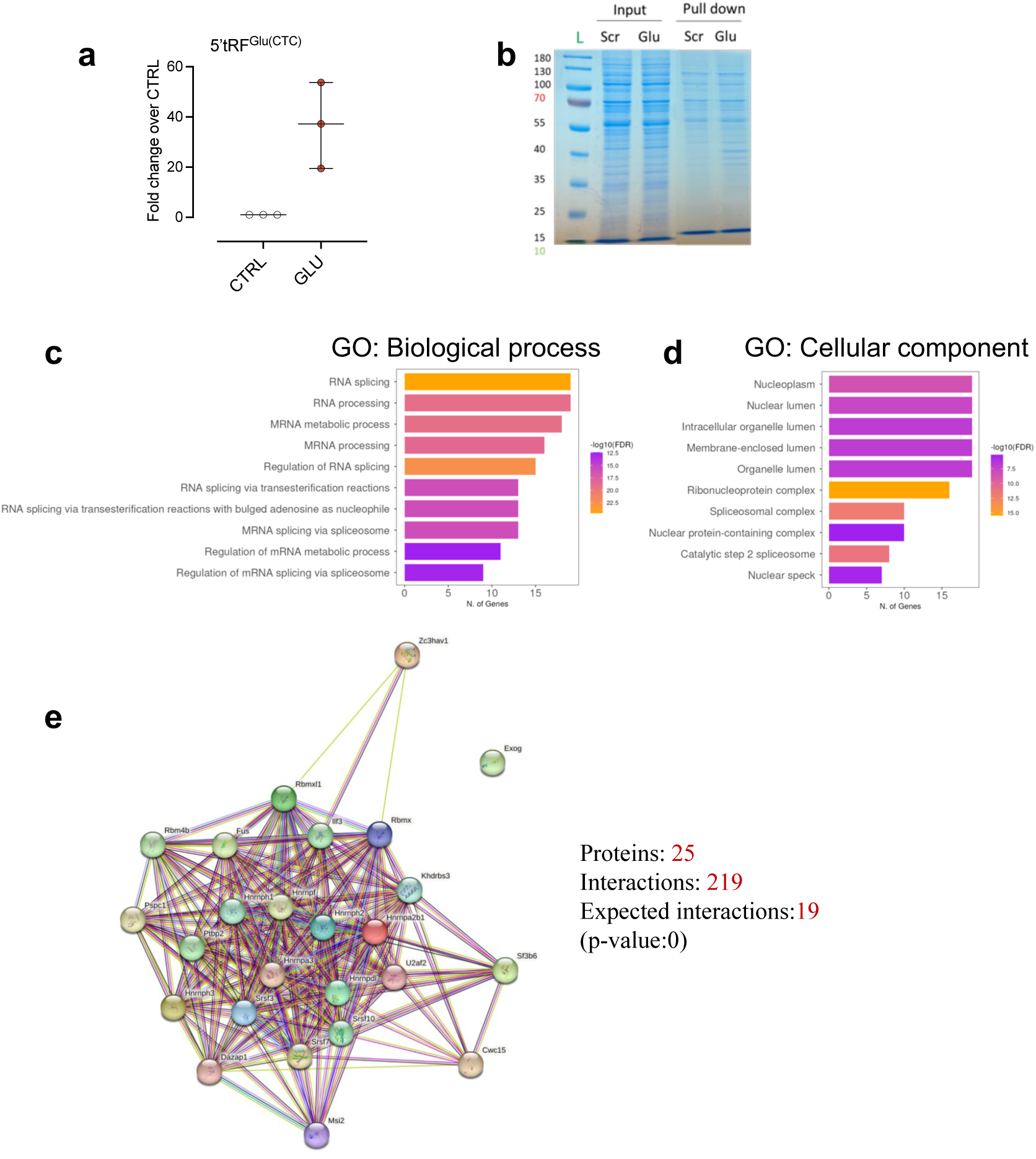
The 5’tRF^Glu(CTC)^ or scrambled (CTRL) biotinylated mimics were transfected in MIN6 *β*-cells; the transfection efficiency was assessed by measuring the levels of 5’tRF^Glu(CTC)^ by qPCR (**a**). Polyacrylamide gel staining show the proteins in the input or eluted samples of pull-down experiment (**b**). **c-d**) Overrepresentation test of GO biological processes (**c**) and GO cellular component (**d**) of proteins pulled-down with GLU mimic. **e**) protein-protein interaction analysis using STRING database of proteins pulled-down with GLU mimic.

**Figure S5:**
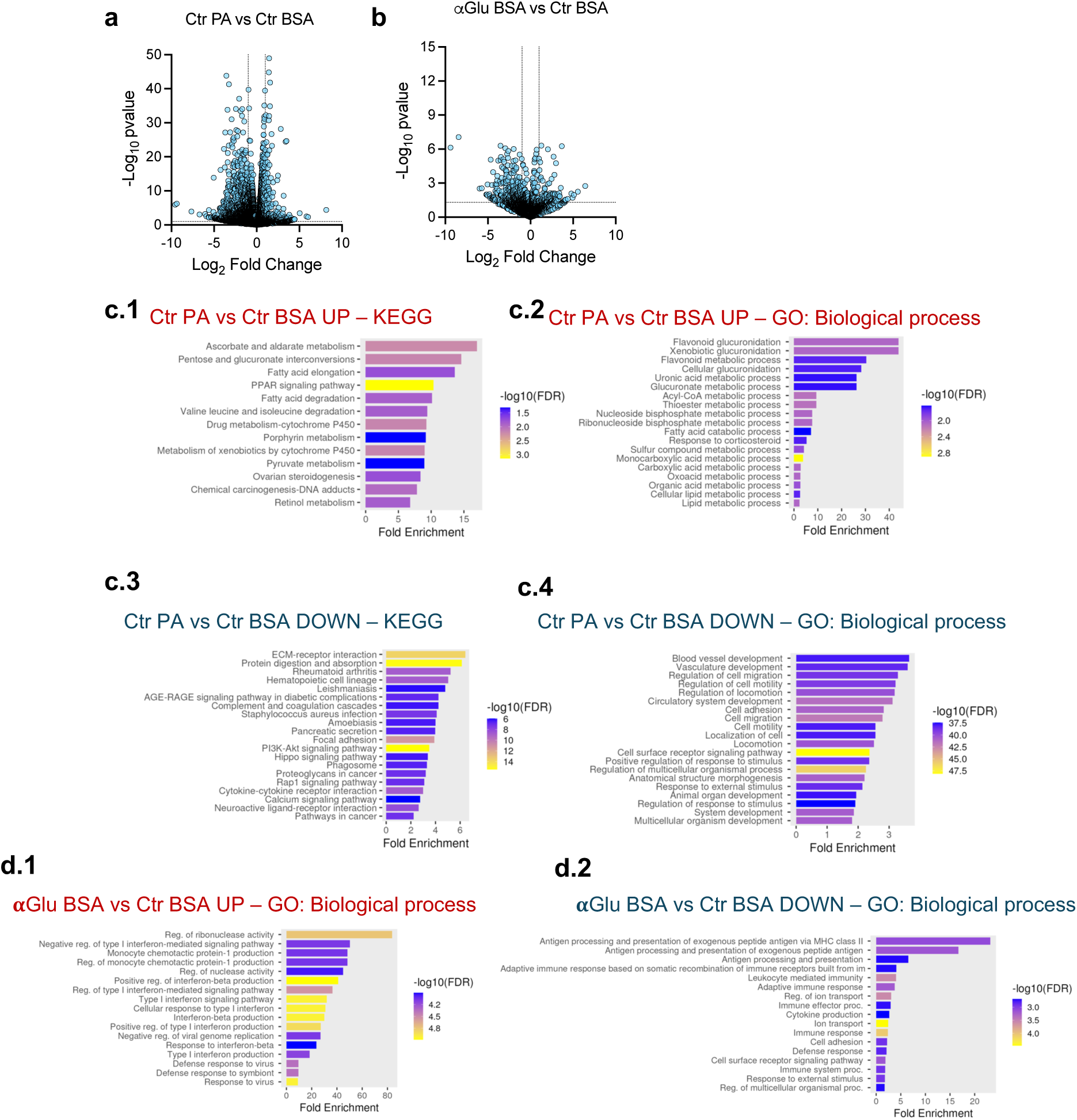
Differential mRNA expression analysis in mouse islets (**a-b**). The volcano plots show the following comparisons: Palmitate (PA) versus BSA condition (**a**; Ctr PA vd Ctr BSA); Transfection with **α**Glu versus transfection with Ctr ASO in basal BSA condition (**b**; **α**Glu BSA vs Ctr BSA). Functional enrichment analysis of gene expression modulation due to PA treatment: bar plots show KEGG and GO biological process terms enriched in upregulated (**c.1-2**) and downregulated genes (**c.3-4**). Functional enrichment analysis of gene expression modulation due to **α**Glu transfection in basal condition: bar plots show GO biological process terms enriched in upregulated (**d.1**) and downregulated (**d.2**) genes.

**Figure S6:**
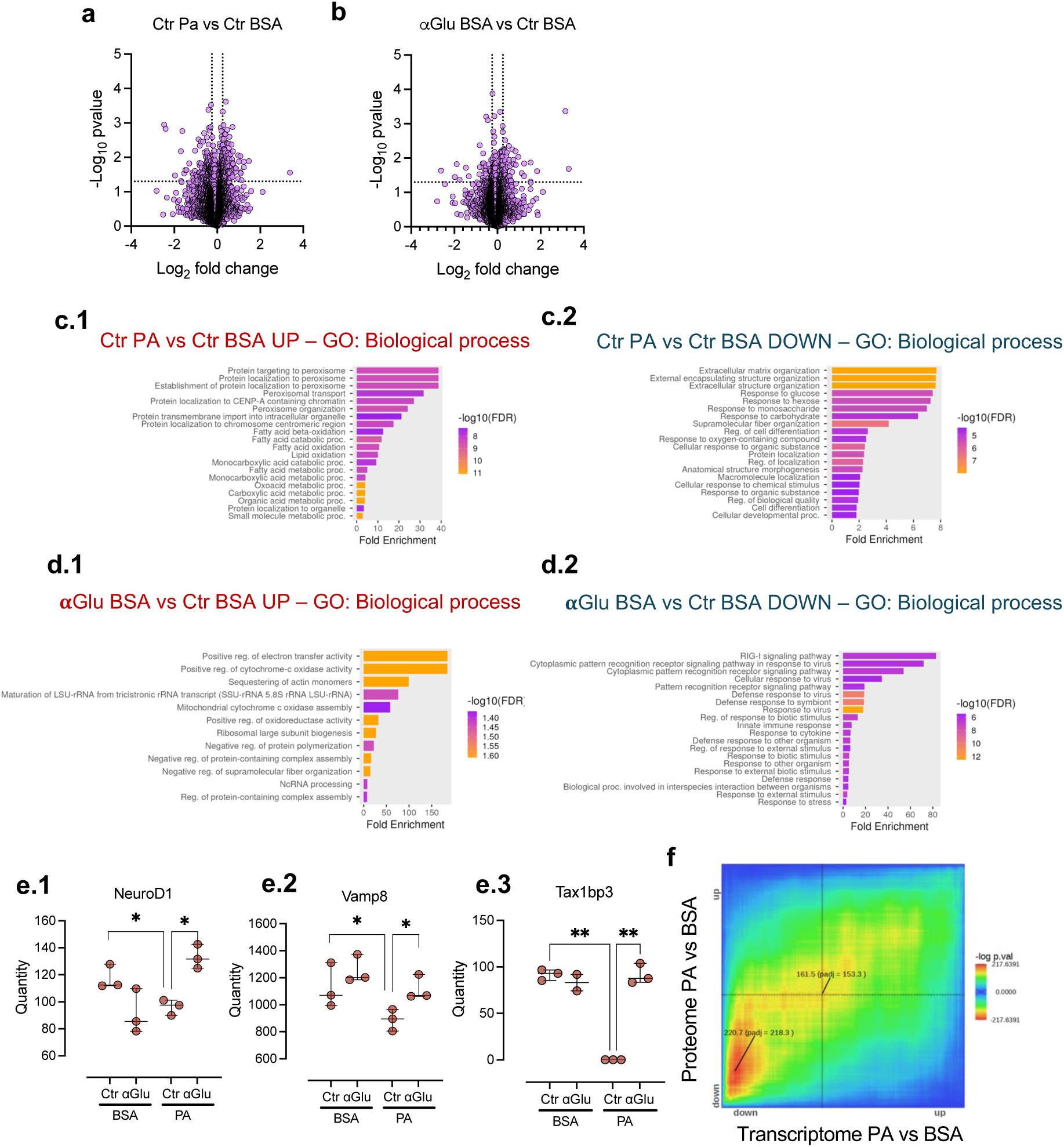
Differential protein expression analysis in mouse islets (**a-b**). The volcano plots show the following comparisons: Palmitate (PA) versus BSA condition (**a**; Ctr PA vd Ctr BSA); Transfection with **α**Glu versus transfection with Ctr ASO in basal BSA condition (**b**; **α**Glu BSA vs Ctr BSA). Functional enrichment analysis of protein modulation due to PA treatment: bar plots show GO biological process terms enriched in upregulated (**c.1**) and downregulated proteins (**c.2**). Functional enrichment analysis of protein modulation due to **α**Glu transfection in basal condition: bar plots show GO biological process terms enriched in upregulated (**d.1**) and downregulated (**d.2**) genes. Quantity values of selected proteins from mass spectrometry data showing the modulation in the different conditions (**e.1-3**). Rank-rank hypergeometric overlap of the modulation in proteome and transcriptome due to palmitate treatment was performed with RedRibbon (**f**).

**Figure S7:**
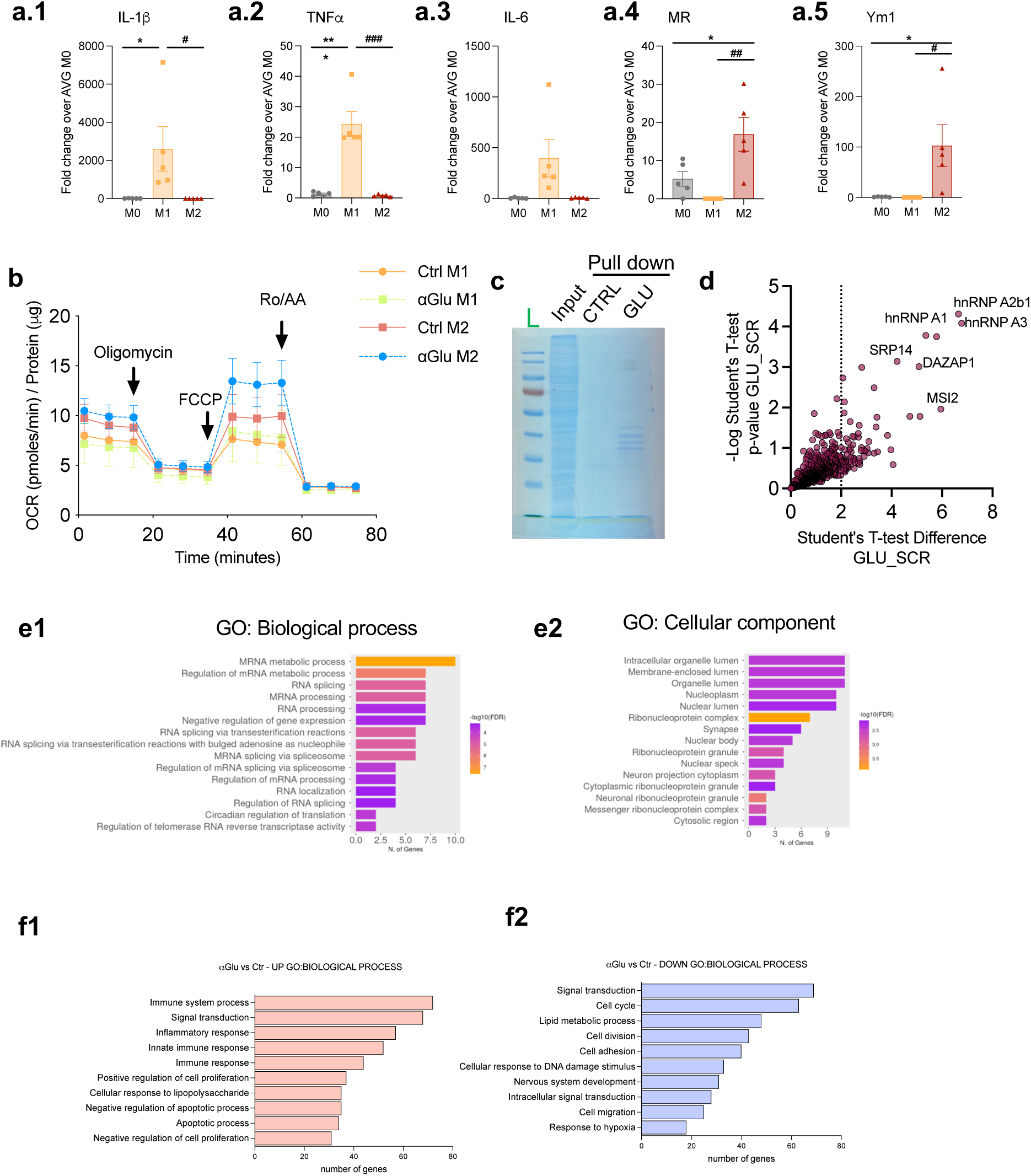
M1 pro-inflammatory and M2 anti-inflammatory polarization was assessed by the expression of gene markers by qPCR (**a1-a5**) OCR was measured in M1 and M2 BMDMs upon transfection with Ctr or *⍺*Glu ASO (**b**). M2-BMDMs lysates were incubated with biotinylated 5’tRF^Glu(CTC)^ mimic (GLU) or with scrambled oligo (CTRL) and then used for pull-down on streptavidin beads. Staining of polyacrylamide gel revealed protein bands present the input and in the eluate of pull-down and from M2 macrophages lysates (**c**). Proteins enriched in the GLU mimic compared to CTRL (**d**) were used for functional enrichment analysis (**e1-e2**). Functional enrichment was performed on upregulated (**f1**) and downregulated (**f2**) genes of M2 macrophages transfected with **α**Glu compared to Ctr.

## Supplementary Tables

**Table S1:** Subjects characteristics and clinical parameters.

| Subject characteristics | NGT (n = 12) | IGT (n = 13) | DM (n = 11) | p-value |
| --- | --- | --- | --- | --- |
| Age (y) | 62.3 ± 2.99 | 65.7 ± 3.74 | 72.7 ± 2.08 | NS |
| Sex assigned at birth (F/M) | 6/6 | 6/7 | 8/3 | -- |
| BMI (kg/m<sup>2</sup>) | 25.5 ± 1.03 | 26.0 ± 1.03 | 26.0 ± 1.11 | NS |
| Fasting glucose (mmol/L) | 5.06 ± 0.16 | 5.23 ± 0.13 | 6.69 ± 0.48 | < 0.001 |
| Medium glucose (mmol/L) | 7.41 ± 0.24 | 8.98 ± 0.41 | 12.7 ± 0.85 | < 0.001 |
| 2-h OGTT Glucose (mmol/L) | 5.86 ± 0.42 | 8.86 ± 0.25 | 15.2 ± 1.19 | < 0.001 |
| Fasting insulin (μUI/ml) | 6.39 ± 0.95 | 9.51 ± 1.34 | 15.0 ± 4.92 | NS |
| Basal ISR (pmol min<sup>-1</sup>m<sup>-2</sup>) | 56.9 ± 6.73 | 82.1 ± 8.46 | 99.4 ± 10.9 | < 0.01 |
| Total ISR (nmol m<sup>-2</sup>) | 40.1 ± 5.00 | 54.5 ± 7.52 | 40.2 ± 4.82 | NS |
| Glucose Sensitivity (pmol min<sup>-1</sup>m<sup>-2</sup>mmol<sup>-1</sup> L) | 86.1 ± 11.8 | 94.1 ± 0.4 | 40.3 ± 8.44 | <0.01 |
| Rate Sensitivity (pmol m<sup>-2</sup>mmol<sup>-1</sup>L) | 1050 ± 290.2 | 656.1 ± 215.1 | 228.3 ± 107.4 | <0.05 |
| Matsuda Index | 7.91 ± 1.45 | 4.06 ± 0.56 | 4.73 ± 1.26 | <0.05 |

**Table S2:**
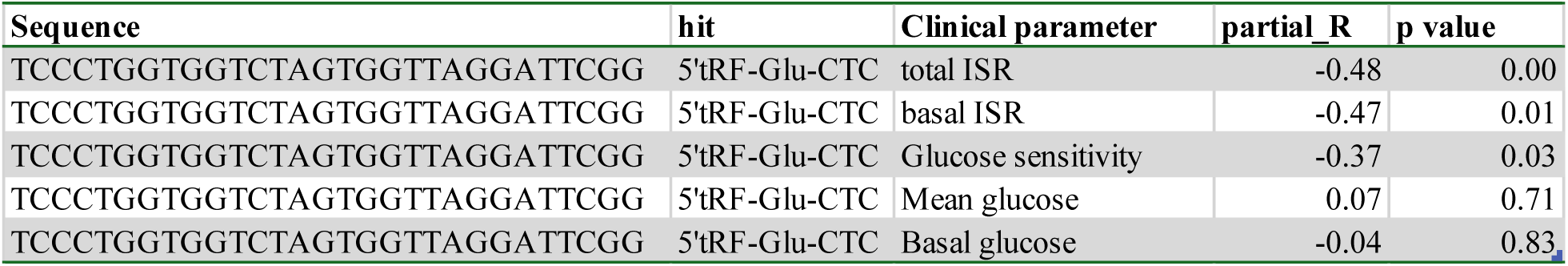
Correlation analysis.

**Table S3:**
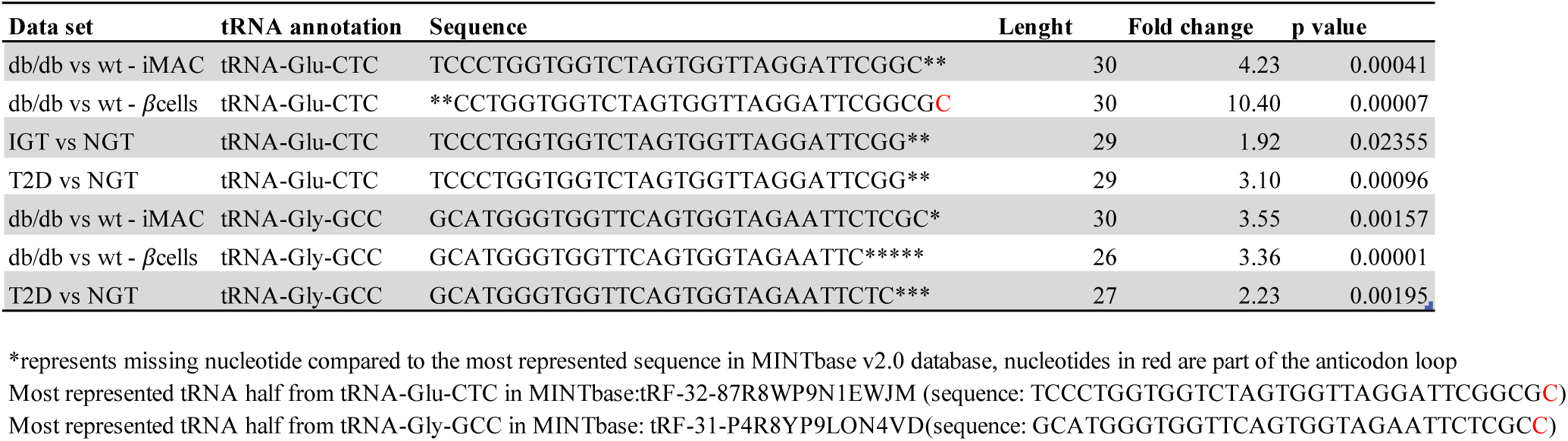
tRF sequences across small RNA sequencing data-sets.

**Table S4:**
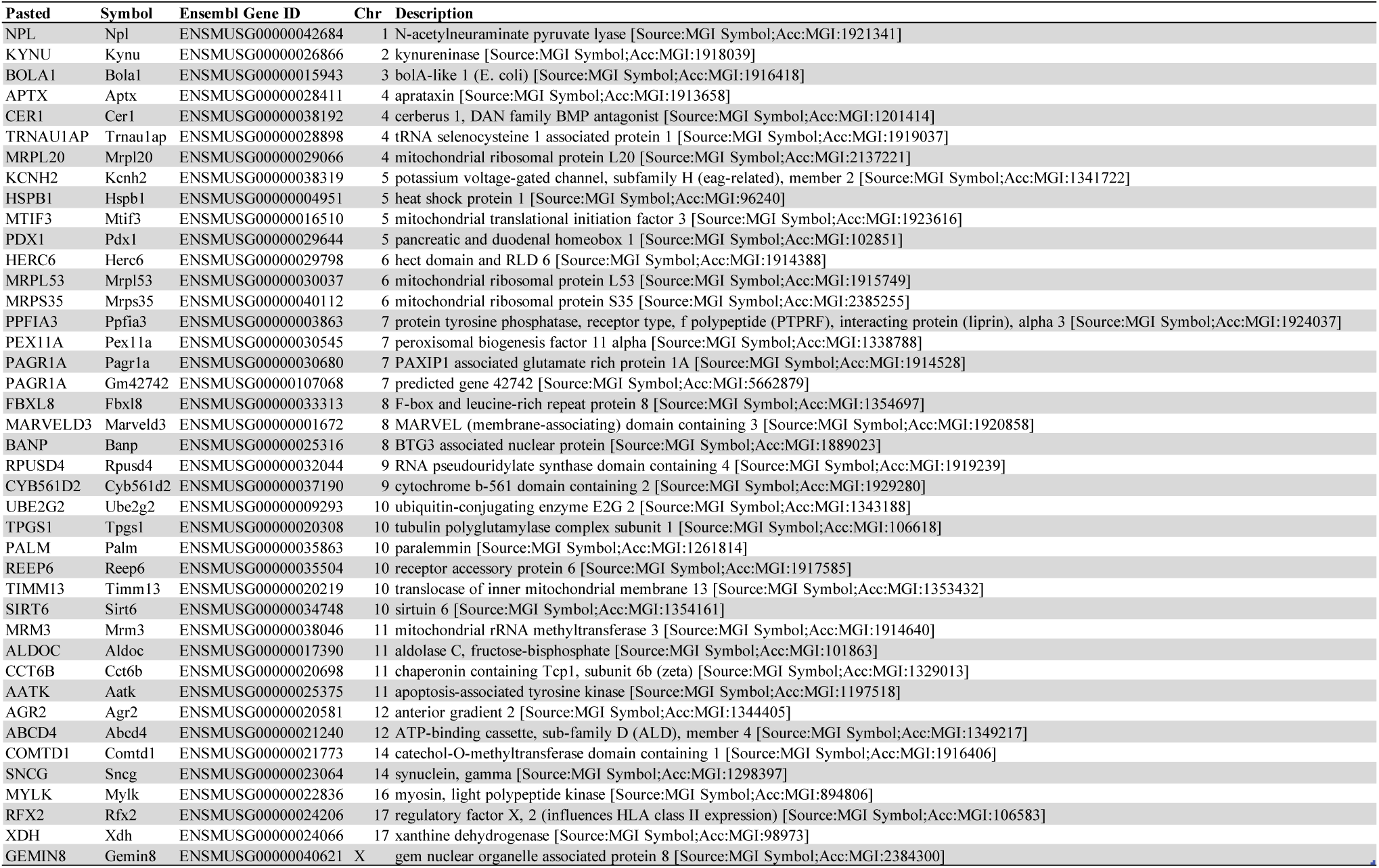
Not-overlapping genes in Rank-rank hypergeometric overlap analysis.

**Table S5:** Human islets from cadaveric donors.

| Preparation | Arrival date | Age | Sex | Weight (Kg) | Height (cm) | Decesse reason | Viability | Purity |
| --- | --- | --- | --- | --- | --- | --- | --- | --- |
| HP2301 | 1/17/23 | 66 | M | 90 | 175 | AVC | 80% | >70% |
| HP2302 | 1/26/23 | 51 | M | 79 | 173 | traumatique non A.V.P | 80-90% | >70% |
| HP2303 | 2/7/23 | 71 | M | 101 | 172 | traumatic | 90% | 90% |
| HP2308 | 6/22/23 | 62 | F | 71.7 | 160 | Rupture d'anévrisme | 90% | >70% |

**Table S6:**
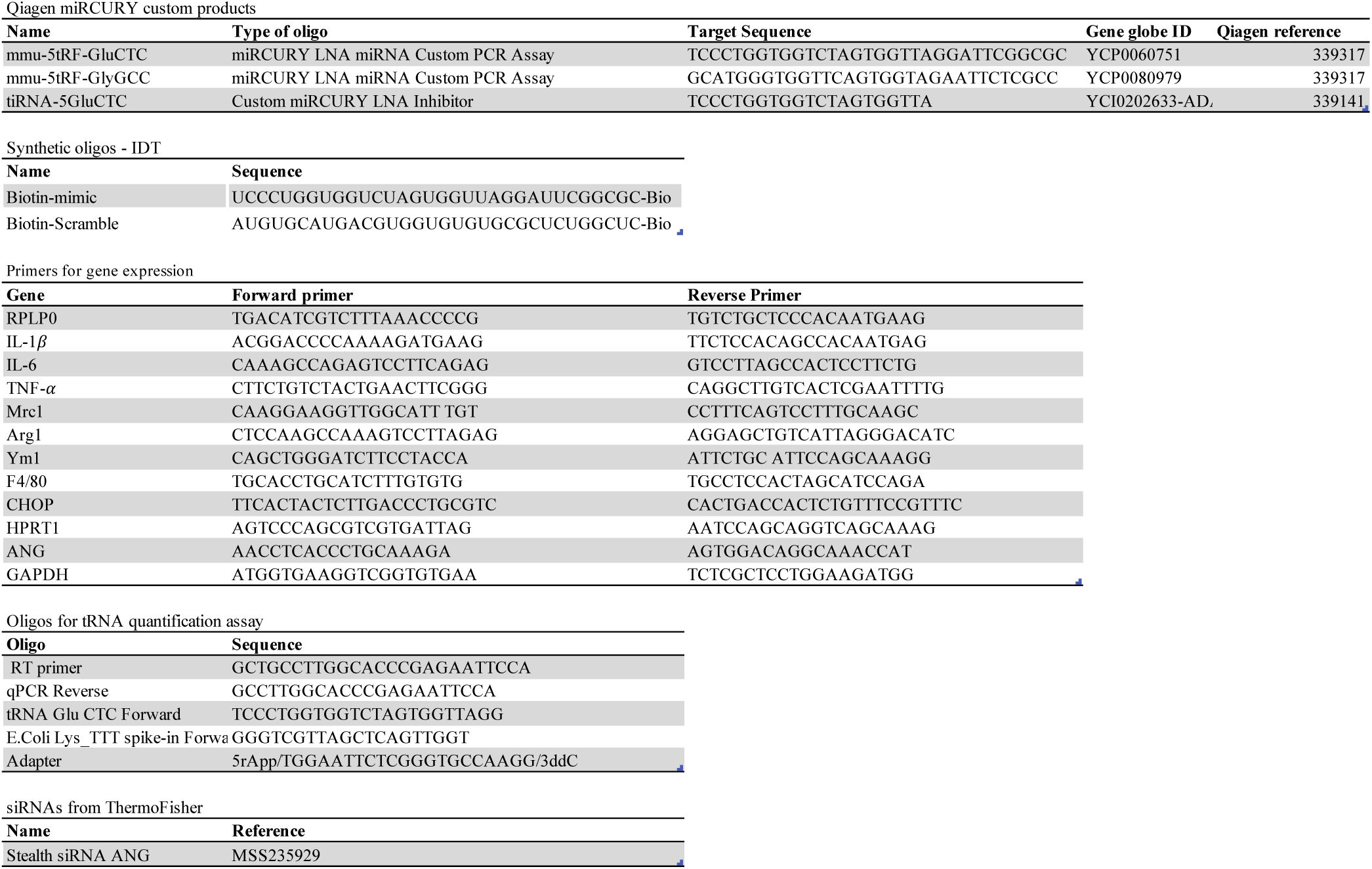
Synthetic oligonucleotides.

**Table S7:** Antibodies and reagents.

| Antibodies |  |  |  |
| --- | --- | --- | --- |
| Name | Company | Reference | Application |
| FITC anti-mouse CD45 Antibody | BioLegend | 103108 | FACS |
| Brilliant Violet 421™ anti-mouse/human CD11b Antibody | BioLegend | 101236 | FACS |
| APC anti-mouse F4/80 Antibody | BioLegend | 123116 | FACS |
| PE anti-mouse CD11c Antibody | BioLegend | 117308 | FACS |
| INS Monoclonal antibody | Proteintech | 66198-1-Ig | Immunofluorescence |
| Cleaved Caspase-3 (Asp175) | Cell Signaling | 9661 | Immunofluorescence |
| MSI2 Polyclonal antibody | Proteintech | 10770-1-AP | Western blot |
| HNRNPA3 Polyclonal antibody | Proteintech | 25142-1-AP | Western blot |
| Reagents |  |  |  |
| Name | Company | Reference |  |
| Oleic acid | Sigma - Aldrich | O1383-1G |  |
| Stearic acid | Sigma - Aldrich | S4751-1G |  |
| Palmitic acid | Sigma - Aldrich | P5585 |  |
| Albumin, Bovine Serum, Fraction V, Fatty Acid Free | Sigma - Aldrich | 126575-10GM |  |
| Murine IL-4 | Peprotech | 214-14 |  |
| Murine IL-13 | Peprotech | 210-13 |  |
| Recombinant Murine IFN-γ | Peprotech | 315-05 |  |
| Murine M-CSF | Peprotech | 315-02 |  |
| Lipopolysaccharid aus Escherichia coli O111:B4 | Sigma - Aldrich | L2630-10MG |  |
| Seahorse XFp Mito Stress Test kit | Agilent | 103010-100 |  |

## Extended material and methods

### RNA pull-down in MIN6 cells

Beads were resuspended in SP3 buffer (2% SDS, 10mM DTT, 50 mM Tris, pH 7.5) and heated 10 min at 75°C to elute proteins. Eluates were digested following the SP3 method (Huges, et al., 2019) using magnetic Sera-Mag Speedbeads (Cytiva 45152105050250, 50 mg/ml). Briefly, samples in SP3 buffer were first treated with 32mM (final) iodoacetamide for 45 min at RT in the dark to alkylate reduced cysteines. Beads were then added at a ratio 10:1 (w:w) to samples, and proteins were precipitated on beads with ethanol (final concentration: 60 %). After 3 washes with 80% ethanol, beads were digested in 50ul of 100 mM ammonium bicarbonate with 1.0 ug of trypsin (Promega #V5113). After 1h of incubation at 37°C, the same amount of trypsin was added to the samples for an additional 1h of incubation. Supernatant were then recovered and transferred to new tubes. Two sample volumes of isopropanol containing 1% TFA were added to the digests, and the samples were desalted on a strong cation exchange (SCX) plate (Oasis MCX; Waters Corp., Milford, MA) by centrifugation to remove traces of SDS. After washing with isopropanol/1%TFA and 2% acetonitrile/0.1% FA, peptides were eluted in 200ul of 40% MeCN, 59% water, 1% (v/v) ammonia, and dried by centrifugal evaporation.

#### LC-MS analysis

Tryptic peptide mixtures were injected on a Vanquish Neo nanoHPLC system interfaced via a nanospray Flex source to a high resolution Orbitrap Exploris 480 mass spectrometer (Thermo Fisher, Bremen, Germany). Peptides were loaded onto a trapping microcolumn PepMap100 C18 (5 mm x 1.0 mm ID, 5 μm, Thermo Fisher) before separation on a C18 custom packed column (75 μm ID × 45 cm, 1.8 μm particles, Reprosil Pur, Dr. Maisch), using a gradient from 2 to 80 % acetonitrile in 0.1 % formic acid for peptide separation at a flow rate of 250 nl/min (total time: 130 min). Full MS survey scans were performed at 120,000 resolution. A data-dependent acquisition method controlled by Xcalibur software (Thermo Fisher Scientific) was used that optimized the number of precursors selected (“top speed”) of charge 2+ to 5+ while maintaining a fixed scan cycle of 2 s. Peptides were fragmented by higher energy collision dissociation (HCD) with a normalized energy of 30 % at 15’000 resolution. The window for precursor isolation was of 1.6 m/z units around the precursor and selected fragments were excluded for 60s from further analysis.

### RNA pull-down in BMDMs

Proteins on beads were digested following a modified version of the iST method (Kulak, et al., 2014) (named miST method). 25 ul of miST lysis buffer (1% Sodium deoxycholate, 100mM Tris pH 8.6, 10 mM DTT), were added to the beads. After mixing and dilution 1:1 (v:v) with H2O, samples were heated 5 min at 75°C. After digestion with 0.5 ug of Trypsin/LysC mix (Promega #V5073) for 1h at 25°C, sample supernatants were transferred in new tubes. Beads were washed with 50 ul of miST buffer diluted 1/1 in H2O, and supernatants pooled with the previous ones. Reduced disulfides were alkylated by adding 25 ul of 160 mM chloroacetamide (32 mM final) and incubating for 45min at 25°C in the dark. Samples were then digested overnight at 25°C with 1.0 ug Trypsin/LysC mix. To remove sodium deoxycholate, two sample volumes of isopropanol containing 1% TFA were added to the digests, and the samples were desalted on a strong cation exchange (SCX) plate (Oasis MCX; Waters Corp., Milford, MA) by centrifugation. After washing with isopropanol/1%TFA, peptides were eluted in 200ul of 80% MeCN, 19% water, 1% (v/v) ammonia, and dried by centrifugal evaporation.

#### LC-MS

Data-dependent LC-MS/MS analyses of samples were carried out on a Fusion Tribrid Orbitrap mass spectrometer (Thermo Fisher Scientific) interfaced through a nano-electrospray ion source to an Ultimate 3000 RSLCnano HPLC system (Dionex). Peptides were separated on a reversed-phase custom packed 45 cm C18 column (75 μm ID, 100Å, Reprosil Pur 1.9 um particles, Dr. Maisch, Germany) with a 4-90% acetonitrile gradient in 0.1% formic acid at a flow rate of 250 nl/min (total time 140 min). Full MS survey scans were performed at 120’000 resolution. A data-dependent acquisition method controlled by Xcalibur software (Thermo Fisher Scientific) was used that optimized the number of precursors selected (“top speed”) of charge 2^+^ to 5^+^ while maintaining a fixed scan cycle of 0.6 s. Peptides were fragmented by higher energy collision dissociation (HCD) with a normalized energy of 32%. The precursor isolation window used was 1.6 Th, and the MS2 scans were done in the ion trap. The *m/z* of fragmented precursors was then dynamically excluded from selection during 60 s.

### LC-MS data annotation and analysis

Data files were analysed with MaxQuant 2.4.7.0 (Cox et al., 2008) incorporating the Andromeda search engine (Cox et al., 2011). Cysteine carbamidomethylation was selected as fixed modification while methionine oxidation and protein N-terminal acetylation were specified as variable modifications. The sequence databases used for searching were the mouse (*Mus musculus*) reference proteome based on the UniProt database (www.uniprot.org, RefProt_Mus_musculus_20230301, containing 55’309 sequences), and a “contaminant” database containing the most usual environmental contaminants and enzymes used for digestion (keratins, trypsin, etc, Frankenfield et al 2022). Mass tolerance was 4.5 ppm on precursors (after recalibration) and 20 ppm on MS/MS fragments. Both peptide and protein identifications were filtered at 1% FDR relative to hits against a decoy database built by reversing protein sequences. All subsequent analyses were done with an in house developed software tool (available on https://github.com/UNIL-PAF/taram-backend/). Contaminant proteins were removed, and iBAQ values (Schwanhäusser et al. 2011) for protein groups were log2-transformed. After assignment to groups, only proteins quantified in at least 4/5 samples of one group were kept. Missing values were imputed based on a normal distribution with a width of 0.3 standard deviations (SD), down-shifted by 1.8 SD relative to the median. Student’s T-tests were carried out among conditions, with Benjamini-Hochberg correction for multiple testing (adjusted p-value threshold <0.05). Imputed values were later removed.

### Proteomic analysis of mouse islets

Cell ysates in RIPA buffer were digested following the SP3 method (Huges, et al., 2019) using magnetic Sera-Mag Speedbeads (Cytiva 45152105050250, 50 mg/ml). Briefly, samples were diluted with SP3 buffer (2% SDS, 10mM DTT, 50 mM Tris, pH 7.5) and heated 10 min at 75°C. Proteins were then alkylated with 32mM (final) iodoacetamide for 45 min at RT in the dark. Beads were added at a ratio 10:1 (w:w) to samples, and proteins were precipitated on beads with ethanol (final concentration: 60 %). After 3 washes with 80% ethanol, beads were digested in 50ul of 100 mM ammonium bicarbonate with 1.0 ug of trypsin (Promega #V5113). After 1h of incubation at 37°C, the same amount of trypsin was added to the samples for an additional 1h of incubation. Supernatant were then recovered and transferred to new tubes. Two sample volumes of isopropanol containing 1% TFA were added to the digests, and the samples were desalted on a strong cation exchange (SCX) plate (Oasis MCX; Waters Corp., Milford, MA) by centrifugation to remove traces of SDS. After washing with isopropanol/1%TFA and 2% acetonitrile/0.1% FA, peptides were eluted in 200ul of 40% MeCN, 59% water, 1% (v/v) ammonia, and dried by centrifugal evaporation.

LC-MS/MS analyses were carried out on a TIMS-TOF Pro (Bruker, Bremen, Germany) mass spectrometer interfaced through a nanospray ion source (“captive spray”) to an EvoSep One liquid chromatography system (EvoSep, Odense, Denmark). Peptides were separated on a reversed-phase Aurora Elite C18 column (15 cm, 75 μm ID, 1.7um, IonOpticks) at a flow rate of 200 nl/min with a 20 sample per day method (runtime time: 68 min, solvents were water and acetonitrile with 0.1% formic acid).

Data-independent acquisition was carried out using a method similar to a standard DIA-PASEF method reported previously (Meier, et al. 2020), with ion accumulation for 100 ms for each the survey MS1 scan and the MS2 scans. Duty cycle was kept at 100%. Precursor ions were chosen within a reduced mobility range from 1/k0 =0.7 to 1.4 and between m/z=400 to 1200. Collision energy was ramped linearly based uniquely on the 1/k0 values from 20 (at 1/k0=0.6) to 59 eV (at 1/k0=1.6). Per cycle, the mass range 400-1200 m/z was covered by a total of 20 windows, each 40 Th wide, with a total cycle time of 1.3 s.

#### Data processing

Identification of peptides directly from DIA data was performed with Spectronaut 19.9 with the Pulsar engine using the “deep” setting and searching the reference mouse proteome (www.uniprot.org) database of February 6^th^, 2025 (54’747 sequences), and a contaminant database containing the most usual environmental contaminants and enzymes used for digestion (from Frankenfield, et al., 2022). For identification, peptides of 7-52 AA length were considered, cleaved with trypsin/P specificity and a maximum of 2 missed cleavages. Carbamidomethylation of cysteine (fixed), methionine oxidation and N-terminal protein acetylation (variable) were the modifications applied. FDR’s for peptide and protein group identifications were all at 1%. Ion mobility for peptides was predicted using a deep neural network and used in scoring. The library created contained overall 228,479 precursors.

Peptide-centric analysis of DIA data was done with Spectronaut 19.9 using the library generated by Pulsar from DIA data. Single hits proteins (defined as matched by one stripped sequence only) were kept in the Spectronaut analysis. Peptide quantitation was based on XIC area, for which a minimum of 1 and a maximum of 3 (the 3 best) precursors were considered for each peptide, from which the mean value was selected. Quantities for protein groups were derived from inter-run peptide ratios based on MaxLFQ algorithm (Cox et al 2014). Global normalization of runs/samples was done based on the median of peptides. Overall 216,168 precursors were quantified in the dataset, mapped to 9’527protein groups. 113,406 precursors (7’584 protein groups) had full profiles, i.e. were quantified in all samples. The average number of data points per peak was 6.1.

#### Data analysis

All subsequent analyses were done with an in house developed software tool (available on https://github.com/UNIL-PAF/taram-backend/). Contaminant proteins were removed, and quantity values form Spectronaut for protein groups were log2-transformed. After assignment to groups, only proteins quantified in at least 3/3 samples of one group were kept. Missing values were imputed based on a normal distribution with a width of 0.3 standard deviations (SD), down-shifted by 1.8 SD relative to the median. Student’s T-tests were carried out among conditions, with Benjamini-Hochberg correction for multiple testing (adjusted p-value threshold <0.05). Imputed values were later removed.

## Notes

### Competing Interest Statement

The authors have declared no competing interest.

